# A cosmopolitan inversion drives seasonal adaptation in overwintering *Drosophila*

**DOI:** 10.1101/2022.12.09.519676

**Authors:** Joaquin C. B. Nunez, Benedict A. Lenhart, Alyssa Bangerter, Connor S. Murray, Yang Yu, Taylor L. Nystrom, Courtney Tern, Priscilla A. Erickson, Alan O. Bergland

## Abstract

Drosophila *melanogaster* living in temperate regions evolve as they track seasonal fluctuations. Yet, we lack an understanding of the genetic architecture of seasonal adaptive tracking. By sequencing orchard populations collected across multiple years, we characterized the genomic signal of seasonal demography and identified that the cosmopolitan inversion In(2L)t drives seasonal adaptation. In(2L)t shows footprints of selection that are inconsistent with simple explanations of genetic drift, as well as signatures of partial selective sweeps. A meta-analysis of phenotypic studies shows that seasonal loci within In(2L)t are associated with behavior, life-history, physiology, and morphology traits. Our results identify candidate regions that underlie seasonal adaptive tracking and link them to phenotype. This work supports the general hypothesis that inversions are important drivers of rapid adaptation.

**One-Sentence Summary:** A chromosomal inversion drives adaptive evolution between seasons in wild fruit flies.

## Introduction

Species living in rapidly fluctuating environments are exposed to temporally and spatially varying selection (*1*). If species harbor polymorphisms that are beneficial in one selective environment but not the other, local adaptation will be evident from shifts in allele frequency across space and time (i.e., adaptive tracking). Context-dependent fitness effects can result in the long-term maintenance of functional genetic variation in populations and even between species (*2*), and can also drive the rapid turnover of new, transiently balanced polymorphisms (*3*).

Recent theoretical work demonstrates that multilocus adaptive tracking is possible (*4*), leaves distinct molecular signatures at linked sites (*5*), and can be facilitated by ecological factors such as seasonal population booms and busts (*6*). Moreover, empirical studies have provided evidence that adaptive tracking can be quantified in both natural and experimental populations (*7*–*9*). Yet, we still have a limited understanding of the ecological drivers that underlie adaptive tracking, its effects on genetic diversity, and its genetic architecture.

Adaptive loci that exist as chromosomal inversions were among the first examples of adaptive tracking (*10*) and are generally thought to be facilitators of adaptation to fluctuating ecosystems (*11, 12*). Given the suppressed recombination across karyotypic states of the inversion, co-adapted alleles existing inside it could be protected from decoupling by recombination (*13*), or from becoming homogenized by gene flow (*14*). Simulations have shown that inversions may capture new beneficial variants that promote local adaptation and should be enriched for loci that are pleiotropic for ecologically important traits (*12, 11*). Empirical work has shown that inversions are often involved in local adaptation among a species’ ecotypes (*15*), show clear correlations to ecological stressors (*16*), and contain alleles in strong linkage disequilibrium with each other, as well as the inversion breakpoints (*17*). Combined, these lines of evidence provide a blueprint to identify adaptive inversions in nature.

Fruit flies (*Drosophila melanogaster*) living in temperate habitats are a premier system to understand the role of inversions in adaptive tracking. Fruit flies have short generation times (∼10-15 days), produce many generations per year (∼15 generations, *18*), and experience fluctuating selection across the changing seasons (*19*). For example, variation in stress tolerance and life-history enable some individuals to better survive the winter months while others more effectively exploit resources in the growing season (*19*–*21*). These observations suggest that seasonal adaptation operates through a resource-allocation trade-off between reproduction and survival that is also mirrored across latitudinal gradients (*22*). Genomic analyses have supported this hypothesis and identified loci whose allele frequencies track the seasons across multiple localities and display parallel changes in allele frequency across spatial gradients (*7, 8, 23*).Analysis of seasonal genomic shifts in Europe and North America identified that the breakpoints of cosmopolitan inversions, particularly of the 10Mb In(2L)t inversion, are enriched for loci that evolve by adaptive tracking (*8*). These findings demonstrate that seasonal adaptation is an intrinsic property of temperate populations and suggest that In(2L)t drives adaptive tracking.

In this paper we test the hypothesis that In(2L)t underlies adaptive tracking across seasonally fluctuating environments. Using genomic time-series data, we asked three questions: 1) Are signals of seasonal selection stronger than temporal drift? 2) What are the ecological drivers of seasonal selection for wild *Drosophila*? 3) Are there signatures of selection and function consistent with the expectations of an adaptive inversion? Overall, our data reveal rapid evolutionary change in response to seasonally varying selection and suggests connections between phenotype, genotype, and the environment. More generally, our work supports the classic hypothesis that inversions are important drivers of adaptation in highly fluctuating environments (*10*).

## Results and Discussion

### Fly populations are structured in both space and time

To characterize patterns of spatial and temporal genetic variation across the temperate range of *D. melanogaster*, we performed principal component analysis (PCA). We focused on localities where flies were sampled at multiple points in time over multiple years from the *Drosophila* Evolution over Space and Time (DEST, *24*) dataset (Munich and Broggingen, Germany; Yesiloz, Türkiye; Odessa, Ukraine; Akaa, Finland; Linvilla [Pennsylvania], Cross Plains [Wisconsin], USA). We also include a densely sampled dataset from Charlottesville (Virginia), USA (37 new pooled samples; **Table S1**). Consistent with previous analyses (*24*), PC1 separates samples from Europe and North America (**Fig. 1A**; Latitude: *F*_*1,86*_ = 586.13, *P* = 3.67×10^−40^; Longitude: *F*_*1,86*_ = 605.47, *P* = 1.08×10^−40^) whereas PC2 separates the eastern and western phylogeographic clusters in Europe (Latitude: *F*_*1,35*_ = 0.019, *P* = 0.89; Longitude: *F*_*1,35*_ = 8.32, *P* = 0.0066). Samples collected at the same locality cluster together, demonstrating that population structure at local scales is stable over time. Yet, these populations also show signals of genetic change from one year to the next (**Table S2**). The overall pattern of year-to-year temporal structure can be visualized through the vector formed among samples collected in subsequent years (**Figs. 1A, 1B**, and **S1**). These shifts in genetic composition from one year to the next suggest that drift is occurring in fly populations and may be especially influenced by seasonal fluctuations in population size.

**Fig. 1:**
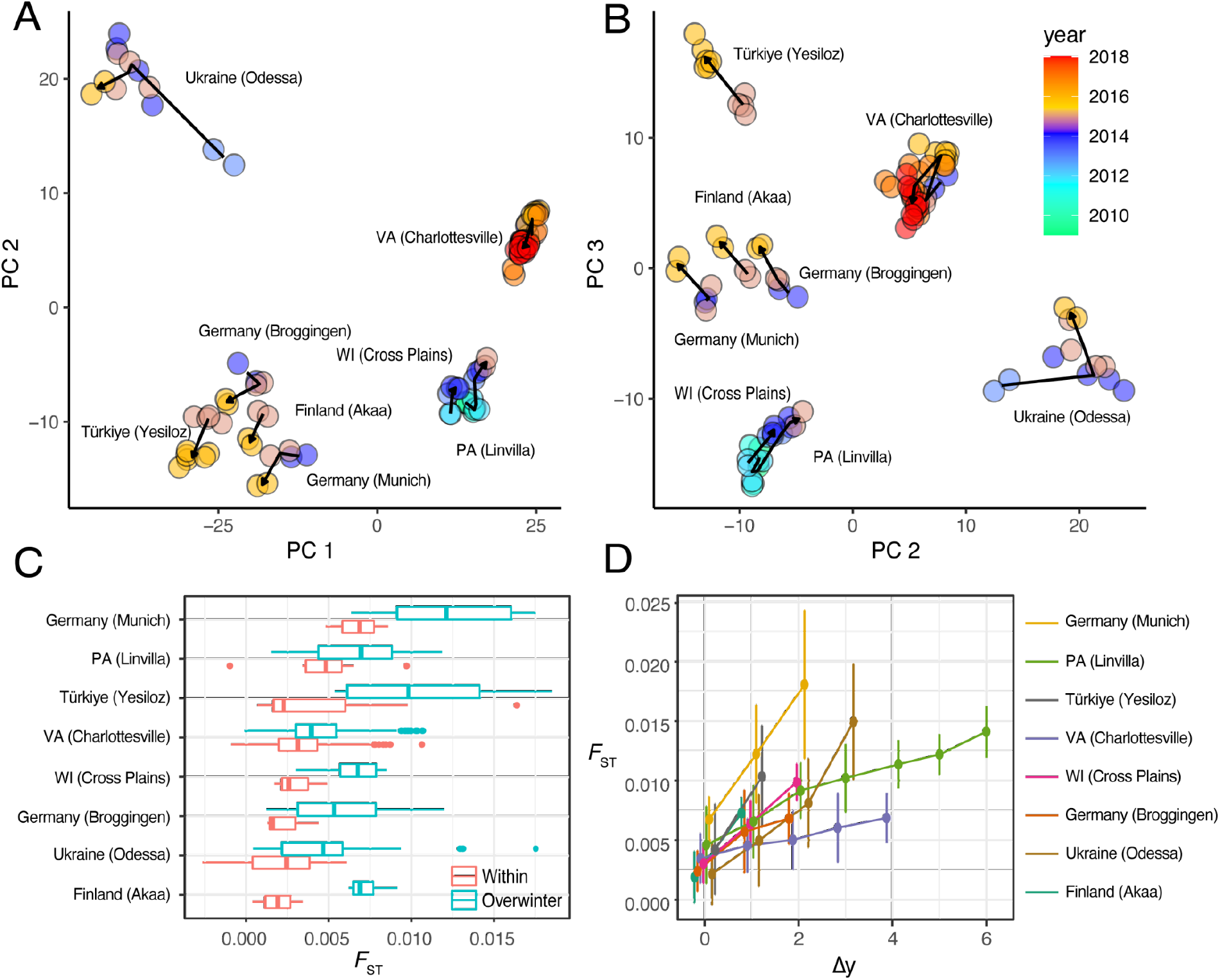
Signatures of overwintering in temperate flies. (**A**) PCA of temporal samples (PCs 1, 2). The arrow path indicates the temporal identity of the samples (arrowheads show the most recent samples, origins show the oldest sample). (**B**) Same as A, but PCs 2 and 3. (**C**) Genome-wide average *F*_ST_ across all within the growing season comparisons (red) and between year comparisons (turquoise). (**D**) *F*_ST_ values across multiple years of collection (Δy is the difference in years of collection; +/-1 standard deviations are shown).

### Temporal structure is driven by seasonal boom-bust demography

To test the hypothesis that seasonal fluctuations in population size influence the genetic composition of populations, we compared patterns of genetic differentiation (*F*_ST_) within a year’s growing season relative to samples collected across years thus reflecting overwintering. Although the exact number of generations in the summer and winter seasons is unknown for fly populations, we presume that the number of generations in the summer must be more than the winter, but that overwintering bottlenecks will nonetheless generate higher levels of differentiation compared to levels of differentiation within a growing season. We observe that the amount of genetic differentiation accrued within the growing season is smaller than that accrued overwinter (**Fig. 1C**; **Table S3**; median *F*_ST-within_ = 0.0031, median *F*_ST-overwinter_ = 0.0045). The pattern of increased differentiation is observed across multiple years of overwintering and the rate of increase of genetic differentiation varies among populations (**Fig. 1D;** ANOVA, *F*_*7,720*_ = 16.72, *P* = 1.60×10^−20^). To quantify the strength of the winter bottleneck, we conducted forward genetic simulations designed to emulate the boom-bust cycle and sampling scheme for the Charlottesville samples (**Fig. S2**). Our results provide support for the hypothesis of yearly population contractions and suggest that the magnitude of winter collapse, in Charlottesville, is on the order of 94% of the maximum summer size (*N*_min_ = 936, *N*_max_ = 16,670, *N*_e_ [effective population size, i.e., harmonic mean of *N*] = 5,599). This inference is consistent with another recent estimate of local effective population size of ∼10,000 (*25*). The signal of year-to-year allele frequency change is distributed across the genome, consistent with a demographic explanation. To show this, we repeated the PCA using random sub-samples of SNPs across autosomes. We then ran correlation analyses of PC projections relative to the year of collection. Our results show that the correlation of PCs 1-3 with year is robust in sample sets larger than 1000 loci (**Fig. S3, Table S4**). Taken together, our analysis suggests that overwintering is a major determinant of the genetic structure of fly populations.

### In(2L)t shows higher temporal F_ST_ than the rest of the genome

Chromosome 2L shows a departure from the temporal signal observed in other chromosomes (**Figs. 2A, S4**, and **Table S4**) suggesting that seasonally varying selection may be acting on the 10Mb cosmopolitan inversion In(2L)t. In principle, the action of seasonal selection on the inversion would lead to an increased temporal *F*_ST_ relative to the rest of the genome where genetic differentiation through time is primarily driven by overwintering drift. To test this hypothesis, we calculated *F*_ST_ within and across years at loci associated with In(2L)t (**Fig. S5**). Mutations inside the inversion in strong association with the breakpoints (Pearson’s correlation *=* 0.95-0.97) show temporal *F*_ST_ values among the top 10-15% relative to matched control distributions (**Fig. 2B**), suggesting that loci in strong association with the inversion are changing in frequency more than expected from overwintering drift. In(2L)t has long been hypothesized to be an adaptive inversion (reviewed in *26*) and our data suggest that seasonality may be an agent of selection at this locus.

**Fig. 2:**
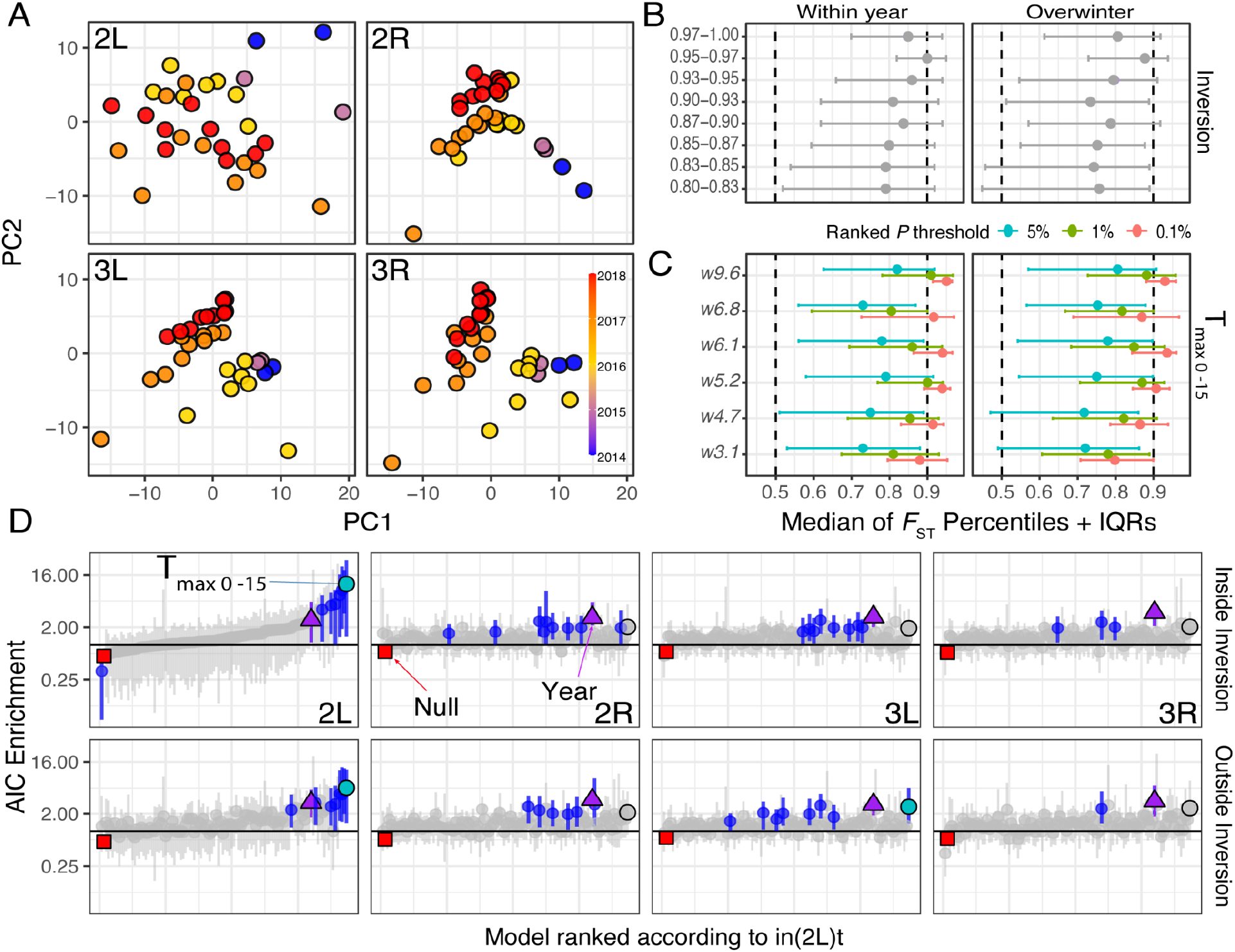
Drivers of temporal structure in Charlottesville flies. (**A**) Chromosome PCAs separated by chromosome. Each point represents one sample. (**B**) *F*_ST_ percentiles from loci associated with In(2L)t breakpoints relative to matched controls. The y-axis shows the correlation coefficient of loci associated with In(2L)t breakpoints. The x-axis shows the percentile of control loci with *F*_ST_ values lower than that of In(2L)t markers. Medians and interquartile ranges (IQRs) are shown. Dashed lines show the 0.5 and 0.9 percentiles. (**C**) Same as B, but for outlier windows in the Charlottesville analysis (see **Fig 3**). (**D**) Environmental models for Charlottesville flies. The x-axis shows models ranked according to the best model in In(2L)t. The y-axis shows the relative rate of enrichment relative to permutations. The vertical line represents the null hypothesis of no change in relative rate between the real and permuted data. Confidence intervals are reported as the 1% and 99% percentiles. Gray circles mean that the confidence intervals contain the value 1. Colored circles represent models whose confidence intervals do not overlap with 1. The model T_max_0-15d, whose confidence intervals do not overlap 1, is indicated in green. The year model is indicated as a purple triangle. The null model (regression against the intercept) is indicated as a red square. The confidence intervals for the null model are smaller than the square and do not overlap with a value of one.

### In(2L)t shows signatures of adaptive tracking and footprints of natural selection

To identify signatures of adaptive tracking, we modeled allele frequency change through time in Charlottesville using a generalized linear model (GLM) at each single nucleotide polymorphism (SNP) in the genome as a function of year of collection and aspects of the environment prior to collection, including temperature, precipitation, and humidity (**Fig. S6**). We then assessed which of these environmental models is found as the best model more often than expected from a permutation analysis. For 2L, both inside and outside of the inversion, the best fit model is the maximum temperature 0-15 days prior to collection (T_max_0-15d). This model is 11 times more likely to be observed as the best model in the real data compared to the permutations (**Fig. 2D**) and shows strong statistical support at top hits (**Fig. S7, Data S1** and **S2**). In contrast to 2L, and consistent with the PCA analysis (**Figs. 2A** and **S3**), changes in allele frequencies on chromosomes 2R, 3L and 3R are best explained by the model that only includes collection year relative to permutations (**Fig. 2D**).

We summarized the output of the T_max_0-15d model using sliding window approaches that test if SNPs whose frequency is strongly correlated with T_max_0-15d are randomly distributed throughout the genome and change with constant magnitude. These analyses indicate that six windows within In(2L)t centered at 3.1, 4.7, 5.2, 6.1, 6.8, and 9.6 Mbs outperform permutations and are enriched for SNPs whose frequency is strongly correlated with recent maximum temperature (**Fig. 3A**). To test whether candidate loci are more differentiated than expected based on short-term demographic fluctuations, we compared temporal *F*_ST_ across candidate SNPs within the windows to matched controls. In Charlottesville, candidate SNPs have higher *F*_ST_ than controls (values fall within the 6% and 10% tails for within and across year comparisons; **Fig. 2C**) thus demonstrating that outlier loci in In(2L)t are strongly differentiated across time. Taken together, the outlier windows identified here represent candidate loci under seasonal selection.

**Fig. 3:**
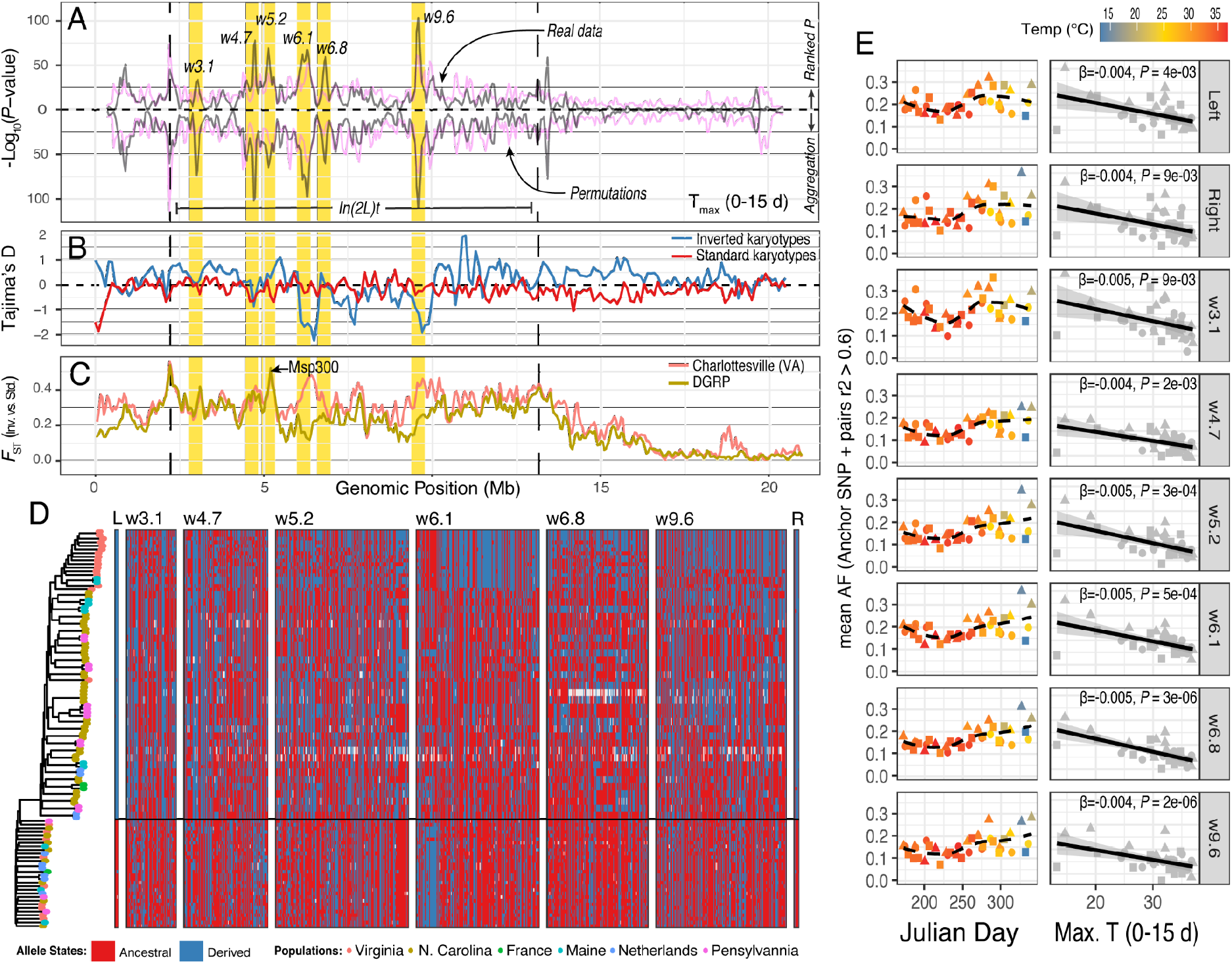
Signals of seasonal selection in In(2L)t. (**A**) Two tests of enrichment for the T_max_0-15d model. The top portion of the panel shows the *P*-value of the rank normalization test. The bottom portion shows the *P*-value of the aggregation test. The pink line shows the 99^th^ quantile of all permutations. The black line shows the real data. The windows of interest are highlighted in yellow. In(2L)t is demarcated by the vertical lines. (**B**) Tajima’s D across 2L for inverted and standard karyotypes. (**C**) *F*_ST_ between the inverted and standard karyotypes in 2L for Charlottesville and the DGRP. (**D**) Haplotype structure plot for inversion breakpoints and windows of interest. Only samples homozygous for the inversion and for the standard karyotype are shown. Samples are sorted and grouped according to the phylogeny (built using candidate SNPs across windows) shown to the left. Ancestral state was determined relative to *D. simulans*. The horizontal line divides inverted vs standard karyotypes. (**E**) Mean allele frequency plots of the anchor loci relative to time (left) and temperature (right). Regression information is shown in each facet. Circles represent samples from 2016, triangles from 2017, and squares from 2018.

These candidate loci show additional footprints of natural selection based on a set of whole genome sequences of individual flies sampled in Charlottesville (see **Table S1**). In(2L)t harbors reduced levels of Tajima’s D at w6.1 and w9.6, consistent with a signature of an incomplete selective sweep (**Figs. 3B, S8A** and **S8B**, *27*). Indeed, w6.1, w6.8, and w9.6 all show low levels of haplotype diversity (**Fig. S9, Table S6**), w6.8 harbors young alleles (**Fig. S8C**), and w9.6 co-localizes with a soft-sweep private to the North American population (*28*). Linkage disequilibrium (LD) analysis across the individual data show strong SNP associations within and among SNPs in our windows of interest (**Fig. S10C**), with the strongest linkage observed between the inversion breakpoints and windows w5.2, w6.1 (**Fig. S10D**). To visualize these patterns of haplotype diversity, we combined our individual Charlottesville data with genome sequence data from inbred lines or haploid embryos established from worldwide collections (see **Tables S6** and **S7**; **Fig. 3D**) and show the presence of soft-sweeps at w6.1 and w9.6. By identifying a series of anchor loci based on LD and GLM scores (**Data S3**) to represent the major haplotypes at each region, we show that the standard karyotype has its lowest frequency in mid-summer and highest frequency in the spring and late fall (**Fig. 3E**; left panels). Regressing the average allele frequency at the anchor SNPs against T_max_0-15d shows a significant negative relationship between In(2L)t haplotype frequency and maximum temperature across 2016-2018 (**Fig. 3E**, right panels).

Estimates of *F*_*ST*_ between standard and inverted karyotypes are highest (*F*_ST_ > 0.4) at the inversion breakpoints and at w5.2 (**Figs. 3C, S10A**, and **S10B**; a signal also observed in the *Drosophila* Genetics Reference Panel [DGRP], *29*). This window primarily encodes the *Msp300* gene, a nesprin-like protein known to mediate the positioning of nuclei and mitochondria in muscle (*30*). Within the region, one seasonal SNP is a trans-species, nonsynonymous polymorphism (**Fig. S11**; 2L:5192177; c.32735G>T, p.Gly10912Val) that is also polymorphic in *D. simulans* and *D. sechellia* (see **Fig. S11D**). While these species are known to hybridize in the wild (*31*), the region corresponding to 32735G>T does not show evidence of introgression between *D. simulans* and *D. sechellia* (*32*). The SNP is in strong linkage with the In(2L)t inversion breakpoints in Charlottesville (mean LD [*r*^2^] = 0.75, sd = 0.03) and present on both karyotypes in at least one African population (**Figs. S11E-S11H**), suggesting that the allele arose prior to both the inversion (75-160 Kya) and the worldwide spread of *D. melanogaster* out of Africa (*33*). Collectively, these findings suggest that In(2L)t captured this old polymorphism during its evolution and is still accumulating beneficial alleles.

### Signals of adaptive tracking within In(2L)t are generalizable to other populations

We tested if the associations between environmental variables and In(2L)t observed in Charlottesville are generalizable to other localities. We used linear modeling to determine the most likely environmental correlates of allele frequency change using temporal samples from localities situated in three distinct phylogeographic regions: Europe west, Europe east, and the east coast of North America (without Charlottesville; see **Fig. S12** and **S13**). The best-fit models for these regions are distinct from those of Charlottesville: average temperature in the 45-75 days prior to sampling for EU-E (T_ave_45-75d; 9.4 times higher than permutations; **Fig. S12A**), average humidity 15-45 days prior for EU-W (H%_ave_15-45d; 5.3 times higher than permutations; **Fig. S12B**), and average temperature 0-7 days prior for NoA-E (T_ave_0-7d; NoA-E does not beat permutations; **Fig. S12C**).

Although the best fit environmental models identified in these other regions are different from what we identified in Charlottesville, the loci underlying allele frequency change in these different regions could be the same. Indeed, the strongest signals of environmental enrichment that we observe in these other phylogeographic clusters is on 2L, and for loci inside In(2L)t (**Fig. S12)**. To test if the same loci change in frequency among these regions, and to test if the direction of allele frequency change is consistent among these regions, we conducted enrichment and directionality tests. The enrichment test asks if top SNPs (top 5%, ranked genome wide) identified in the candidate windows in Charlottesville are also in the top 5% of SNPs identified in the other regions. The directionality test asks whether the direction of allele frequency change between best-fit GLM models for each group are the same, conditional on SNPs being in the top 5% for both groups. The directionality test is calculated as the proportion of SNPs with the same sign of allele frequency change with respect to the environmental variable that we identify as the best fit on 2L (**Fig. S12**). The null hypothesis is 50%, and values significantly different from 50% in either direction indicate that there is some consistency in direction. Candidate loci at w3.1, w5.2, w9.6 and the inversion breakpoints are enriched for SNPs strongly correlated with weather in both Charlottesville and either EU-E or EU-W (Fisher’s exact test, *P* < 0.05; **Fig. 4A**). We observe signals of under-enrichment at w6.1 and w6.8 when contrasting Charlottesville to EU-W and EU-E, suggesting that the incomplete sweeps observed in this region may be private to North American populations. The directionality test shows that nearly all top SNPs are changing in consistent directions in Charlottesville and EU-E and EU-W (directionality scores >90% or less than 10%; binomial test, *P* < 0.05, for most candidate loci; **Fig. 4B**). The enrichment and directionality tests show that the loci that we identify in Charlottesville are fluctuating in other locations, and that the SNPs at these candidate loci are changing in frequency as a haplotype block, consistent with fluctuations of a large linked locus.

**Fig. 4:**
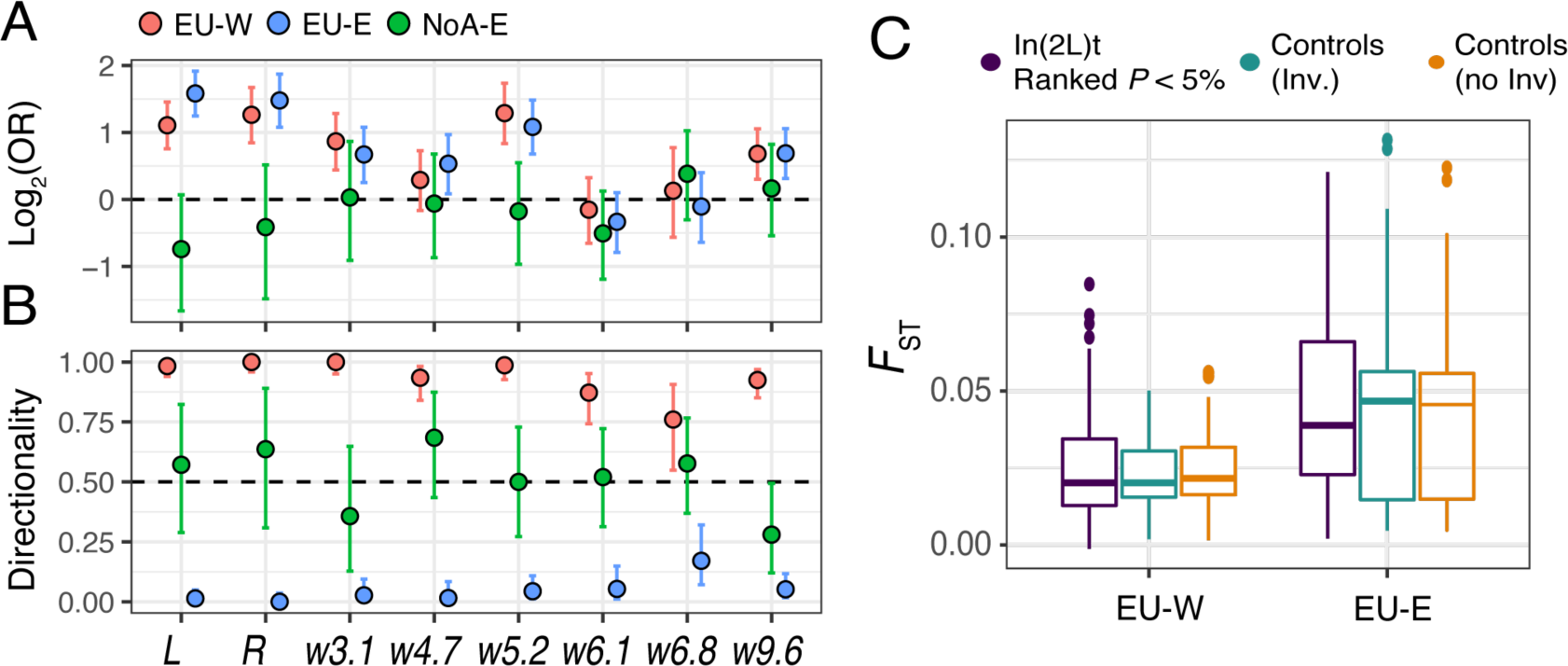
Enrichment, directionality and genetic differentiation in In(2L)t. (**A**) Enrichment scores (odds ratio [OR]) between loci in the T_max_0-15d model in Charlottesville and the best models across other populations at inversion breakpoints and windows in In(2Lt). 95% confidence intervals shown. The horizontal line is the null expectation of no enrichment. (**B**) Same as E but for directionality scores. The horizontal line is the null hypothesis of no consistent directionality. (**C**) Spatial *F*_ST_ for three types of markers: candidate SNPs in In(2L)t (ranked *P* < 5% in their respective best models), and controls inside and outside inversions in EU-W and EU-E. The comparisons were done within the EU-W and EU-E clusters separately.

### Seasonal drivers of allele frequency change at In(2L)t

The specific aspects of weather identified by our analysis vary by geographical region (T_max_0-15d in Charlottesville, T_ave_45-75d in EU-E, H%_ave_15-45d in EU-W, and T_ave_0-7d in NoA-E). Are these aspects of weather the proximate causes of temporally varying selection, or do they reflect something else? We consider three non-mutually exclusive hypotheses. First, allele frequencies oscillate across seasons as a direct consequence of fluctuating environmental selection. Although the specific environmental models that we identify suggest different stressors drive selection across the species range, these variables may simply be proxies for a shared seasonal stressor. Second, due to the temporal nature of our data it is plausible that allele frequencies are driven by negative frequency dependent selection (*34*), and because weather is seasonal, artefactual associations with environmental variables may have emerged. A third hypothesis is the joint action of genetic overdominance and boom-bust demography. In this model, the inverted and standard karyotypes are maintained via heterotic (*35*) or associative overdominance (*36*) and are kicked out of equilibrium by yearly bottlenecks. As selection returns alleles back to equilibrium frequency, allele trajectories may resemble seasonal oscillations.

Although we cannot rule out any of these models conclusively, our data are most in-line with the seasonal stressor hypotheses. To arrive at this conclusion, we first consider the overdominance-perturbation model. Under this model, we predict low spatial differentiation and lower than average temporal differentiation because natural selection would be rapidly pushing populations back to a common equilibrium. To the contrary, our data show high temporal differentiation within Charlottesville (**Fig. 2C**), yet only average differentiation across spatial gradients (Kruskal-Wallis tests; *P*_EU-E_ = 0.47, *P*_EU-E_ = 0.46; **Fig. 4C**). Although evidence in favor of the seasonal stressor model over the negative frequency dependent selection model is more limited, and differentiating these hypotheses is challenging (*34*), several pieces of evidence point in favor of the seasonal stressor model. The first is a comparison between our results and several previous studies. In one, seasonal frequency change of In(2L)t was documented during the 1980’s in a Spanish population (*37*). There, In(2L)t is high frequency in the fall and low frequency in the summer, similar to what we observe (**Fig. S14**). In(2L)t was also found to be higher frequency in the fall compared to the summer in some North American populations (*38*), and time series analysis of caged seasonally evolving populations shows that In(2L)t increases frequency from summer to fall (*9*). The seasonal stressor hypothesis is interesting to consider in light of spatial patterns of allele frequency change at In(2L)t. In *D. melanogaster*, inversions are generally thought of as “hot adapted” (*39*), and most cosmopolitan inversions are higher frequency in more tropical locales across multiple continents (*38*). However, In(2L)t is peculiar in this regard. It shows a stable latitudinal cline only in Australia and weak or unstable patterns of clinality in other continents (*26*). In(2L)t is found at frequencies between 20-80% in many high latitude locales (**Table S1**), in contrast to other cosmopolitan inversions (*26*). Taken together, patterns of spatial and temporal allele frequency change show that In(2L)t is common across the range, weakly differentiated across spatial gradients, and highly differentiated through time, suggesting that In(2L)t frequencies are affected by temporally heterogeneous selection.

### In(2l)t loci are associated with ecologically important traits

Although individual candidate loci underlying seasonal evolution in *D. melanogaster* have been identified and validated (e.g., *40, 41*) genome-wide analysis of seasonal allele frequency change in this species has provided limited resolution to identify phenotypes and loci underlying adaptive tracking (*7, 8*). To elucidate the phenotypic consequences of the candidate loci that we identify, we aggregated line mean estimates for 225 phenotypes collected by dozens of labs using the DGRP (**Tables S8** and **S9**). Phenotypic variation of 36 traits is correlated with In(2L)t inversion status, and these traits span all phenotypic categories (**Figs. 5A** and **S15A**). We performed GWAS for each trait and assessed the level of enrichment between loci that affect traits and loci that are strongly associated with the T_max_0-15d model in Charlottesville. We show that In(2L)t is enriched for loci that are associated with phenotypic variation in the DGRP and are also strongly correlated with T_max_0-15d in Charlottesville (**Figs. 5B** and **S15B**). We also investigated the proportion of SNPs that have the same sign of allele frequency change conditional on those SNPs being in the top 5% of both the GWAS and the GLM models (i.e., “directionality”; see **Fig. 4B**). For each SNP under investigation, we used the estimated allelic effect from the GWAS and the slope of allele frequency change with respect to the T_max_0-15d model. Like our previous directionality analysis, our null hypothesis is 50%. Values significantly different from 50% show evidence of consistent alignment of effect directions between the GWAS and GLM analysis.

**Fig. 5:**
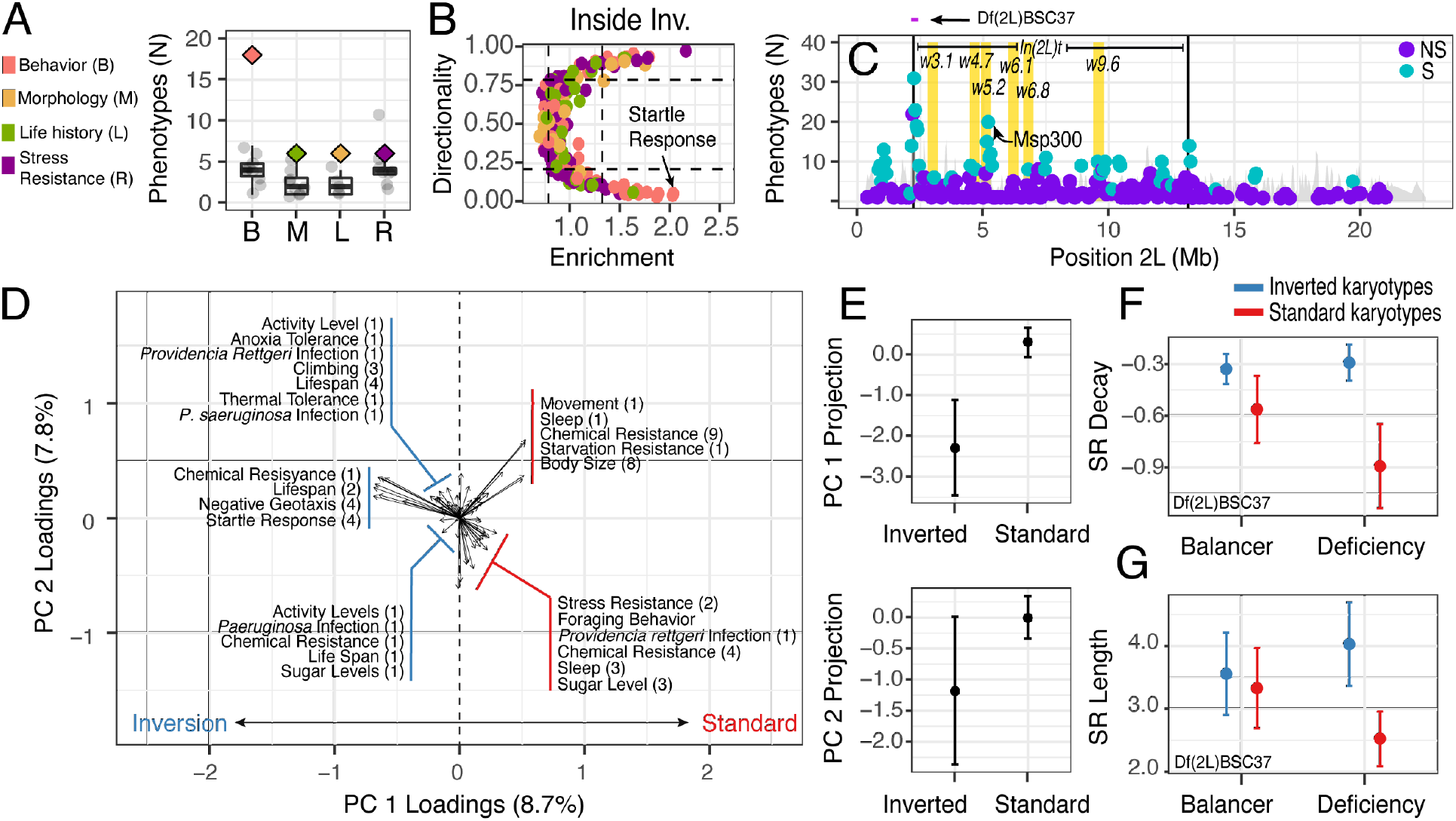
Phenotypes associated with candidate loci on chromosome arm 2L. (**A**) Number of GWAS phenotypes associated with inversion status in the DGRP. Traits are divided across four phenotypic categories. The real data are shown as diamonds, permutations are shown as black points and boxplots. (**B**) Directionality and enrichment analysis between the DGRP-GWAS and the best environmental model in Charlottesville. Lines indicate null expectations estimated by permutations. Each point is a phenotype and colors same as A. (**C**) Window level enrichment analysis across 2L. The y-axis shows the window-wise number of enriched phenotypes (i.e., significant in both the GLM and GWAS). Windows that exceed permutation are shown in turquoise, otherwise in purple. (**D**) PCA of phenotypes associated with the candidate SNPs in the inversion. Each arrow represents a phenotype characterized in a GWAS study. Number of studies are in parentheses. (**E**) Inversion status is significantly associated with PC1 but not PC2. The 95% confidence intervals are shown. (**F**) Quantitative complementation tests using deficiencies shows that the inverted and standard karyotype have significantly different effects in the deficiency background but not the balancer background for the decay rate of startle response. (**G**) Same as F but for the startle response length.

SNPs on 2L that are associated with phenotypes and T_max_0-15d show levels of directionality greater than we expect from permutations, a signal only seen in 2L (**Figs. 5B** and **S15B**). We performed the enrichment analysis on sliding windows across 2L to test for localized enrichment of SNPs that are top hits for both the GWAS and GLM relative to permutations. Windows near the inversion breakpoints and at w5.2 show enrichment of GWAS and GLM hits relative to permutations and are associated with many phenotypes. Regions near the left breakpoint are associated with at least 31 phenotypes, whereas w5.2 is associated with at least 23 phenotypes (**Figs. 5C** and **S15C**). The association between these loci and phenotypic variation in the DGRP may have been missed in previous studies because inversion status is explicitly used as a cofactor in GWAS (*42*).

The inverted and standard alleles impact a suite of traits, demonstrating pleiotropy. To characterize patterns of trait covariation, we conducted PCA using all phenotypes identified in our sliding window analysis (**Fig. 5D**). Presence of the inversion is significantly associated with PC1 (*t-test, t* = -4.1, df = 19.8, *P* = 0.00042; **Fig. 5E**), but not PC2 (*t* = -1.3, df = 20.3, *P* = 0.18) or PC3 (*t* = -1.2, df = 22.0, *P* = 0.22). Inversion homozygotes show higher levels of basal and induced activity, lifespan, and resistance to a variety of stressors whereas standard homozygotes show higher values for starvation resistance and chemical resistance. To validate the phenotypic effect of allelic variation at the candidate regions, we focused on startle response. Startle response is a top hit in our association analyses (**Fig. 5B**), and inverted homozygotes have a greater startle response than standard homozygotes. We show that the inverted haplotype is higher frequency in spring and late winter compared to the summer and fall (**Fig. 3E)** and suggest that startle response is important for overwintering survival and recolonization because it could increase the chance of finding shelter during the winter and patch recolonization during the summer.

We used quantitative complementation to validate the effect of candidate windows on startle response. We crossed selected DGRP lines to 5 deficiency bearing lines for regions in In(2L)t (see **Table S10; Fig. S16**). One deficiency that covers the left-inversion breakpoint (2.17-2.45 Mb; **Fig. 5C**, top) fails to complement the inverted and standard alleles for two measures of startle response: the rate of return to basal activity (**Fig. 5F**; *χ*^2^_SR decay_[df = 3] = 24.20, *P* = 2.26×10^−5^) and the startle-response length (**Fig. 5G**; *χ*^2^_SR length_[df = 1] = 3.504, *P* = 0.061; see **Table S11**). Complementation tests confirm that the inversion increases startle response, consistent with the direction of effect among inbred DGRP lines. Our analysis and validation of the phenotypic effect of inversion associated candidate loci underlying seasonal adaptive tracking therefore generates hypotheses for future studies and suggests, for the first time, that behavioral traits underlie seasonal adaptation.

### The role of adaptive inversions in fluctuating environments

Inversions have been proposed as drivers of adaptation for populations living in fluctuating environments because they can protect co-adapted alleles from being broken up by recombination or swamped by migration (*11*). Yet evidence for the ecological drivers of the phenotypes implicated in adaptation is scarce. Here, we have shown that In(2L)t is an adaptive locus that facilitates seasonal evolution in *Drosophila* and provides links between phenotype and genotype. Overall, this work supports the general hypothesis that inversions are key facilitators of adaptation to fluctuating ecosystems.

## Materials and Methods

### Fieldwork

Fly collections were completed at a local orchard in Charlottesville, VA (Carter Mountain, 37.99N, 78.47W) from 2016 to 2019. Collections from 2016 to 2018 were done using aspirators and netting every two weeks starting in mid-June when peaches come into season in central VA and ending in mid-December at the end of the fall season. The collection in 2019 was done at the beginning of the growing season in June. Because *D. melanogaster* is phenotypically similar to its sister taxa *D. simulans*, we determined species identity using the male offspring produced from isofemale lines set from wild-caught flies. *D. melanogaster* isofemale offspring were frozen in ethanol and stored at -20°C prior to sequencing.

### DNA extraction, sample preparation and sequencing

We prepared two sets of samples: Pool-seq samples, and individual DNA-seq libraries. For pool-seq, we prepared 37 libraries (sample-size per pool are available at **Table S1**). Libraries were made from individual male flies collected from 2016, 2018, and 2019. For 2016, we prepared 119 individual samples collected across various time points of the year’s growing season. For 2018, we prepared libraries for 43 individuals collected in the fall. For 2019, we prepared libraries for 41 individuals collected in the spring. For the pooled samples, DNA extractions were done using the extraction protocol outlined in (*7*). Extracted DNA was diluted in a 1:1 DNA:water ratio mixture, then sonically sheared to create fragments 500 bp in length using a Covaris machine. Library preparation was done with a NEBNext Ultra II kit following the kit’s protocol. Eight cycles of PCR were done in the final PCR enrichment step. Following library preparation, each pool was quantified and pooled in equal concentration to produce a final sequencing library. This final library was size-selected on a Pippen for DNA sizes in the 600-750 bp range and sequenced on a NovaSeq using 2×150 paired-end (PE) reads. For individual libraries, DNA extractions were done using an Agencourt DNAdvance kit (Beckman-Coulter A48705). Library preparation was done using a diluted Nextera kit (Illumina FC-131-1024) (*43*) using a liquid handling robot. Unique index sequences were generated according to (*44*). Sequencing of the 2016 individuals was done on an Illumina HiSeqX (2×150). Sequencing for the 2018 and 2019 individuals was done on an Illumina Novaseq (2×150).

### Pooled sequences - Integration into to the DEST dataset

Quality control, mapping, SNP calling and dataset merging was done using the DEST dataset mapping pipeline (https://github.com/DEST-bio/DEST_freeze1) using the optimized settings for the PoolSNP caller (*45*) and enforcing a global average minimum allele frequency of 1%. The DEST mapping pipeline accounts for potential contamination with *D. simulans* in the pools using competitive mapping (*8, 24*). We combined the Charlottesville pool-seq with the pool-seq samples from DEST to generate a new dataset that contains 283 pooled samples from 22 countries across 12 years 2003-2018. This version of the dataset contains 3,265,012 SNPs. We further filtered pools to include only samples that had more than two consecutive years of sampling and were sampled least twice within a year. The resulting data set is composed of samples from six countries: Finland (n=6 pools), Germany (n=12), Spain (n=6), Türkiye (n=14), Ukraine (n=19), and the USA (n=63). These can be further subdivided into 14 localities: Akaa (Finland), Broggingen and Munich (Germany), Yesiloz (Türkiye), Odessa (Ukraine), Charlottesville (Carters Mountain Orchard in Charlottesville, VA, USA), Cross Plains (WI, USA), and Linvilla (Linvilla Orchard in Media PA, USA). Repetitive elements, defined by the Interrupted Repeats, Microsatellite, RepeatMasker, SimpleRepeats, and WM_SDust tracks from UCSC Genome Browser (http://genome.ucsc.edu) were removed from further analysis.

### Individual sequences - DNA mapping, QC, and phasing

Prior to mapping all individual PE reads were merged into longer reads using bbmerge (*46*). Merged reads were trimmed using bbduk v38.98, flags: *ftl*=15 *ftr*=285 *qtrim*=w *trimq*=20. Reads were mapped to the *Drosophila* genome (Release 6 plus ISO1 MT; https://www.ncbi.nlm.nih.gov/assembly/GCF_000001215.4/) using BWA-MEM v0.7.17 (*47*). Bam file sorting and read deduplication was done using Picard tools v2.27.4 (https://broadinstitute.github.io/picard/). Bam file quality was assessed with qualimap v2.2.1 (*48*). In the case of samples that were sequenced across multiple lanes, bam files were joined into a single-individual bam using samtools v1.9 merge (*49*) prior to PCR duplicate removal. GVCFs were created using the HaplotypeCaller program of the GATK v4.2 pipeline (*50*). SNP calling was done by first generating a GenomicsDBI object using GATK’s GenomicsDBImport. SNP calling was done using the GenotypeGVCFs program. SNPs were calibrated using VariantRecalibrator using the DGRP as the training set. We used WhatsHap v1.7 (*51*) to conduct read-based phasing, followed by population based phasing using shapeit v4.2.2 (*52*). For the individual based sequencing, we identified 6,689,236 autosomal SNPs that passed filtering. Repetitive elements, defined by the Interrupted Repeats, Microsatellite, RepeatMasker, SimpleRepeats, and WM_SDust tracks from UCSC Genome Browser (*53*) were removed from further analysis.

### Other fly datasets used

*D. melanogaster* data was downloaded from three public repositories: DEST (*24*), DGRP2 (*26*), and DPGP3 (*54, 55*). *D. simulans* was obtained as a VCF file from Zenodo’s repository of (*56*). Data for *D. yakuba* was obtained from (*57*) and mapped to its corresponding genome (NCBI acc. GCA_016746365.2). Data for *D. sechellia* was obtained from (*32*) and mapped to its corresponding genome (NCBI acc. GCF_004382195.1). Data for *D. mauritiana* was obtained from (*58*) and mapped to its corresponding genome (NCBI acc. GCA_004382145.1).

### Temporal analysis using PCA

Principal component analyses (PCA) were conducted on the Pool-seq time series data from the DEST dataset as well as the new Charlottesville samples. These analyses were done on two ways: First using global data using populations from Munich and Broggingen, Germany; Yesiloz, Türkiye; Odessa, Ukraine; and Akaa, Finland: Linvilla, Pennsylvania, and Cross Plains, Wisconsin, USA, including all Virginia (Charlottesville, Virginia, USA) samples. This is the PCA presented in **Fig. 1**). The second way only used the data from Virginia (shown in **Fig. 2**). In both cases, we applied a minimum allele frequency filter of 0.001 across the subset of populations. We also applied a mean effective coverage *N*_eff_ filter of 28 (see explanation below). *N*_eff_ was calculated as in (*59*):

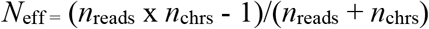

Where *n*_reads_ is the read depth, and *n*_chrs_ is the number of pooled chromosomes. *N*_eff_ is calculated in a SNP-wise manner, and the mean *N*_eff for_ a given sample is used in our filtering. We also applied local missing data filters of 0.01 (i.e., any samples with higher missing data is excluded). The resulting dataset (a matrix of allele frequencies) was used to calculate the principal component analysis using FactoMiner v2.6 (*60*). The *N*_eff_ filter of 28 was determined empirically by running the PCA analysis sequentially at various *N*_eff_ thresholds. We observe that when samples with *N*_eff_ < 28 are included in the analyses these samples create outliers in PCA driven by *N*_eff._ When PCA is done with samples *N*_eff_ > 28, *N*_eff_ no longer influences clustering across major PCs.

In addition, we randomly sampled SNPs in increments of 100, from 100 to 1000 SNPs, and in increments of 1000, from 1000 to 20000 SNPs and performed PCA, and calculated correlations of PC 1, 2, and 3 with year of collection (ρ^2^_year_), frequency of inv(2L)t (ρ^2^_inv(2L)t_), and effective coverage (ρ^2^_*Neff*_). In parallel, we ran an identical analysis but with sample labels permuted. We repeated this process 500 times each and compared estimates of the real order of the data relative to permutations.

### Generalized linear models (time and environment)

We tested the effects of time and temperature on allele frequencies (AFs) by fitting a binomial generalized linear model (GLMs) using *fastglm* v0.0.3 (*61*). We used the effective coverage *N*_eff_ as the observed sample size for the GLMs. For each SNP, we fit 112 models. First, we fit a null model in which AFs were regressed onto their means (AF = β_0_ + *ε*;) and a second model where AFs were regressed onto the collection year as an unordered factor (AF = β_0_ + β_1_(y_factor_) + *ε*). Next, we constructed 110 environmental models (AF = β_0_ + β_1_(y_factor_) + β_2_(γ) + *ε*; where γ is an environmental covariate). For any model, the environmental covariate is a summarization (e.g., mean) of temperature, precipitation, or humidity in the weeks to months prior to sampling. For temperature, precipitation, or humidity, we summarized hourly estimates from the NASA-power dataset (*62*). using the mean and variance over the selected window of time. For temperature, we calculated the maximum and minimum hourly temperature in the selected window of time. For temperature, we also calculated the number of days where the daily maximum was above 32°C or minimum daily temperature was below 5 °C. See **Fig. S6** for a visual depiction of the summarization scheme.

We performed likelihood ratio tests (LRT) between each environmental model and the year-only model; we performed LRT between the year-only model and the null model (**Data S1**). For other DEST populations we fit similar models, but the environmental models have the form AF = β_0_ + β_1_(Population:y_factor_) + β_2_(γ) + *ε*, a change also included in the time model (AF = β_0_ + β_1_(Population:y_factor_) + *ε*).

For each SNP, and for the four population sets (Charlottesville, EU-E, EU-W, and NoA-E) independently, we identify the “best” model by assessing the number of times were the model fit in the real data is the best fit model (by AIC) relative to 100 permutations (i.e., the model’s relative rate of enrichment). Notice that the permutations were done preserving linkage relationships in the data, meaning that each SNP was permuted in the same way and thus the correlation of signal across that is generated by linkage is preserved in each permutation.

Window based enrichment analyses were done in two ways. First, by rank-normalizing the LRT *P*-values such that all *P*-values are transformed into a uniform distribution bounded between 1 and 1/L. By using these rank-normalized *P*-values and dividing the genome into 10 Kb windows with a 5 Kb step, we assessed the number of SNPs in each window that had *P*-values at the 5% of the ranked *P-value* distribution. We calculated a p-value of enrichment for each window under the null hypothesis that 5% of SNPs in the window will be in the top 5%, ranked genome-wide using the binomial.test() function in R. We also calculated the *wZa* metric, which is based on Stouffer’s method (*63*) for aggregating *P-values* across windows, and we use it to assess the strength of signal across the genome.

### Inference of inversion 2Lt markers

We used the DGRP (*26*) to identify a panel of SNPs associated with inversion breakpoints for Inv(2L)t using only lines karyotyped as standard or inverted homozygotes; no heterozygous lines were used (*42*). We conducted a PCA on the DGRP panel using only data from chromosome 2L (**Fig. S6**). We identified putative inversion markers using PCA loadings followed by estimating levels of linkage disequilibrium (LD) using Plink v1.9 (*64*). We only kept SNPs with highest loadings to PC 1 and mean LD > 0.99 relative to the inversion kayrotype. We validated these putative inversion SNPs by calculating LD of these markers in our individual phased data from Charlottesville (SNP positions were transformed to Dmel 6 using LiftOver; https://genome.ucsc.edu/cgi-bin/hgLiftOver). We kept 47 SNPs with mean LD > 0.8 in our Charlottesville data as a list of final inversion markers. We trained a linear support vector machine model (SVM) using the 47 markers and the DGRP data using the R package “e1071” v1.7-11 and used this SVM to perform *in-silico* karyotyping of the sequenced inbred lines.

### Forward genetic demographic simulations

To test if variable bottleneck sizes influence patterns of genetic differentiation through time, and to infer minimum and maximum population sizes during boom-bust cycles that are consistent with our data we performed genetic simulations. First, we performed a coalescent based neutral simulation of a single population with θ_π_ = 0.001 using *msprime* (*65*) in python 3.8. This neutral background was used as a burn-in within the forward genetics software, SLiM3 (*66*). SLiM3 was used to simulate cyclic population crashes genetic data, and we varied the population size maximum (nMax) and the population size minimum (nMin) under a model of instantaneous change in population size. For each parameter combination, the simulated demographic event had a constant population size at nMax from generations 1-16, 19-33, and 36-50 and the bottlenecks occurred at generations 17-18 and 34-35 where the population size was set to nMin. A VCF of 50 simulated diploid individuals was output at the end of each generation to track allele frequency changes. Allele frequencies were simulated to mimic pooled-sequencing using the sample.allele() function in the poolSeq v0.3.5 (*67*) package with a mean coverage of 60. Principal component analyses were performed using the PCA function in FactoMiner v2.6 (*60*) and missing data was imputed using the mean. Pairwise *F*_ST_ was calculated using the compute.pairwiseFST() function in poolfstat v2.1.1 (*68*). Every parameter combination was simulated 100 independent times with different seeds. Model selection was performed using approximate bayesian computation (ABC) under a rejection method with a threshold of 10% using *abc* v2.1(*69*) in R. The summary statistics used were the medians of: within year *F*_ST_, between year *F*_ST_, median allele frequency variance, the *r*^2^ of PC1, PC2, PC3, LD1, and LD2 values with simulation year (**Fig. S2**).

### Cross-model enrichment and directionality scores

We tested whether candidate loci that show strong signal in Charlottesville are enriched for those SNPs in the top 5% identified in the best models in EU-W, EU-E, and NoA-E using Fisher’s exact test. We also assessed if allele frequency changes are consistent between population sets by calculating the proportion of SNPs that have the same sign of allele frequency change with respect to the population cluster’s best fit model, conditional on their being strong allele frequency change at all (top 5% in both population clusters). Because the specific environmental models for EU-W, EU-E, and NoA-E are different from those identified in Charlottesville, directionality values of either 0% or 100% indicate that alleles at a candidate window are changing in frequency as a haplotype block relative to Virginia. To this end, we performed a binomial test on the windows of interest across the best models in EU-W, EU-E, and NoA-E relative to Charlottesville. The significance of a window’s directionality is assessed if the estimate of the binomial test is different from 50% (i.e., random expectation).

### Population genetic analyses

For our phased samples, we calculated *F*_ST_, π, Tajima’s D, and haplotype numbers in vcftools v0.1.16 (*70*). LD was calculated using plink v1.9 (*64*). TMRCA was calculated using GEVA v1.0 (*71*). For pool-seq data, *F*_*ST*_ was calculated using poolfstat v2.1.1 (*68*). Temporal *F*_*ST*_ was calculated among populations sampled across timepoints. Spatial *F*_*ST*_, in Europe, was done on samples collected during the fall of 2015 to ensure temporal homogeneity across comparisons. For the haplotype-inferred trajectory analyses, anchor markers were selected based on *Q*-values < 0.05 in the pooled data (i.e., adjusted *P*-values using false discovery rate). Using the LD estimates from the individual data, we identified all loci pairs (+/-0.2 Mb) with r^2^ > 0.6 in Virginia to the anchor locus. We used the averaged frequency of the anchor loci and its high LD pairs as estimators of haplotype frequencies in the pooled data (Anchor loci and LD pairs can be found in **Data S3**).

### Matched controls

Matched controls, used for *F*_ST_ analyses, were identified by sampling the genome for 100 SNPs with similar recombination rate (+/-0.20 cm/Mb), global allele frequency (+/-0.030). By design matched controls must originate from chromosomes different from the one containing the SNP of interest.

### Phenotypic association with inversion status

GWAS meta-analyses was performed using phenotypic datasets described in **Table S8**. We constructed a phenotypic dataset from published data of DGRP line averages for 225 phenotypes (**Table S9**). We annotated these phenotypes by classifying each phenotype into one of four general-groups: “Behavior”, “Life-History”, “Morphology”, and “Stress-resistance”. We used this dataset to establish the effect of cosmopolitan inversions (In(2L)t, In(2R)Ns, In(3L)P, In(3R)K, In(3R)Payne, In(3R)Mo) on phenotype using a linear models designating inversion presence focusing on DGRP strains reported to be homozygous inverted, or homozygous standard. In the case of 3R the analysis was implemented to identify traits associated with any inversion inside the chromosome.

### GWAS analysis

We performed GWAS for each phenotype using the DGRP2 dataset with GMMAT v1.3.2 (*72*). In this analysis our null model is described by the formula: Phenotype_i_ = β_0_ + β_1_(Wolbachia) + β_2_(GRM), where β_2_(GRM) is a random effect matrix. The null model is compared to a full model defined as: Phenotype_i_ = β_0_ + β_1_(Wolbachia) + β_2_(SNP dosage) + β_3_(GRM). Where β_1_(Wolbachia) is a fixed effect corresponding to Wolbachia infection status of the line as defined by DGRP2, and β_3_(GRM) is a random effect matrix. The GRM refers to the genetic relatedness matrix. The GRM is traditionally used to account for cryptic relatedness or genetic structure among the individuals or strains. The publicly available GRM generally used in DGRP-GWAS has large sections around the inversions removed but despite this remaining relatedness within the matrix is influenced by inversion status of the lines, especially In(2l)t (*42*). As the goal of our analysis was to reveal the effect of elements within inversion on these phenotypes, our analysis used an identity matrix in place of the standard GRM.

### GWAS-GLM enrichment & directionality

We identified regions of the genome that are enriched for SNPs identified in the GWAS analysis and loci identified by the T_max_0-15d model from Virginia. We grouped our SNPs per chromosome and inversion status (inside or outside), then tabulated the number of SNPs that are outliers for the GLM and GWAS datasets (top 5% SNPs, ranked by p-values), and calculated odds-ratios using the Fisher’s Exact Test. We calculated the directionality score as the proportion of times the sign of allele frequency change was identical for the GLM and GWAS models, using SNPs in the top 5% for both. A direction score of 1 thus indicates that the sign of allele frequency change at every SNP under investigation is the same, and 0 indicates that all SNPs have opposing signs of effect in the GLM and GWAS analysis. We repeated this analysis with the same 100 GLM permutations referenced previously to develop an empirical null distribution for the enrichment and directionality tests.

After using the previous analysis to identify phenotypes broadly enriched for seasonal SNPs, we performed a sliding-window analysis to identify specific regions of high enrichment. The sliding analysis used a window size of 100 kb and a step of 50 kb and examined the local enrichment of phenotypes with SNPs that are in the top 5% of the GLM and GWAS hits, per phenotype. After identifying the phenotypes enriched with GLM and GWAS at each window. We replicated this sliding window analysis using the 100 GLM permutations, described above. We identified significant windows as those were the Fisher’s exact test p-value of enrichment per phenotype had chromosome-wide Bonferroni-corrected *P*-value < 0.05, and where the number of phenotypes that had such a significant enrichment was greater than 100% of the permutations. This analysis identified 62 phenotypes as candidate phenotypes. We scaled these phenotypes to conduct a principal component analysis using FactoMiner v2.6 (*60*).

### Startle response quantitative complementation tests

We performed quantitative complementation using deficiencies to test the effect of In(2L)t on startle response by selecting a set of 5 deficiency lines covering regions within our windows of interest. We additionally selected DGRP lines that included lines both homozygous for inverted and standard karyotype and included lines with either the ancestral or derived haplotypes for w5.2 and w9.6. We used these deficiency lines, and constructed a set of 25 F1 crosses (see **Table S10**: Crossing Scheme). For example, the Df(2L)BSC37, dpp[EP2232]/CyO deficiency (Bloomington #7144) which spans approx. 2.1 mb to 2.5 mb was crossed with three inverted DGRP lines and two standard. For each F1 cross, we sorted 3–5 day old females into balancer and deficiency F1 backgrounds based on the curly wings balancer phenotype.

To test startle response, we mounted Trikinetic monitors (DAM2 *Drosophila* Activity Monitor) to a vibrating pad (Best Choice Products #SKY3197) within a Percival incubator held at 20°C in constant light. On five consecutive days we assayed a new set of flies from each cross to avoid a day-effect, and on each day randomly selected the location of each fly amongst the Trikinetic monitors and wells to avoid a monitor-effect. We placed the flies in the monitors overnight with DAM (v3.10.7) software set to record at 1 second intervals. At 10 am the following morning we set the vibrating pad to the lowest setting for 5 seconds, before letting the monitors continue data collection for at least 10 minutes. We repeated the whole experiment in three independent blocks. Prior to data analysis, we smoothed the activity data for each fly as activity counts over a window size of 90 seconds, progressing with an interval of 30 seconds.

We estimated the startle response in two ways. First, we calculated startle duration as the time between a fly’s peak activity post-stimulus, and their return to that flies basal activity level as defined as the average activity prior to startle. We tested for a failure to complement using a mixed effect model implemented in *lme4* version v1.1-30 (*73*), by performing a likelihood-ratio test between the full model to the additive one:

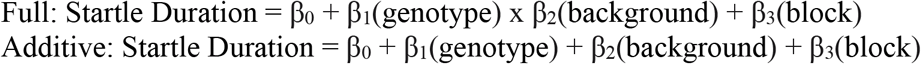

Where, genotype is a fixed effect that includes both the inversion status and the ancestral/derived state of any haplotype in that window, background is a fixed effect factor that considered if that fly has the balancer or deficiency background, and block is a random effect factor that considers which of the three experimental blocks the data was collected from.

The second way that we assessed startle response was by estimating the rate of change of startle induced activity following stimulation. The startle-response decay rate is the slope of activity per unit time following stimulation. We scaled activity by dividing the activity rate scores by the basal activity rate, defined as the average activity per minute over the hour prior to stimulus, for each individual fly before fitting models to these scaled activity scores over the span of time starting at peak activity post-stimulus. The rate of change of activity following startle is the “startle response decay” and we estimated these decay rates using the Emmeans v1.8.1-1 (*74*). To test for failure to complement using a mixed effect model, we conducted a likelihood-ratio test of the full model to the additive one:

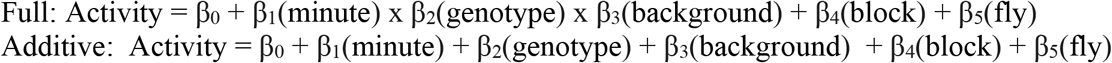

Where β_1_(minute) is a fixed effect that tracks the elapsed time in minutes since the peak of post-stimulus activity, β_2_(genotype) is a fixed effect that includes both the inversion status and the ancestral/derived state of any haplotype in that window, β_3_(background) is a fixed effect factor that considered where that fly has the balancer or deficiency background, β_4_(block) is a random effect factor that considers which of the three experimental blocks the data was collected from, and β_5_(fly) is the random effect of each individual fly.

## Acknowledgements

The authors wish to acknowledge members of the DrosEU consortium for their discussion and insightful feedback related to this paper. To L. Galloway, A. Perrier, H. Makowski, K. Lamb, A. Lopez, and J. Jiranek at UVA for editorial comments during early drafts of the manuscript. To the staff of the Carter’s Mountain Orchard and the owners of Chiles Family Orchards for allowing us to visit their orchard to sample fruit flies. The authors acknowledge Research Computing at The University of Virginia for providing computational resources and technical support that have contributed to the results reported within this publication. URL: https://rc.virginia.edu.

## Author contributions

Conceptualization: JCBN, AB, PAE, AOB

Methodology: JCBN, BAL, AB, CSM, PAE, AOB

Investigation: JCBN, BAL, AB, CSM, YY, TLN, CT, PAE, AOB

Visualization: JCBN, BAL, CSM, AOB Funding acquisition: AOB

Project administration: JCBN, AB, AOB Supervision: JCBN, PAE, AOB

Writing – original draft: JCBN, BAL, AB, AOB

Writing – review & editing: JCBN, BAL, AB, CSM, YY, TLN, PAE, AOB

## Ethics Statements

All collections were done either on public lands, or in private properties with the express consent of the owners. The authors declare no competing interests.

## Data Availability

This paper used multiple datasets that are available in the National Center for Biotechnology Information (NCBI; https://www.ncbi.nlm.nih.gov), as described in their corresponding citations. Data generated as part of this paper is also available in NCBI in bioprojects: PRJNA882135, PRJNA728438, and PRJNA727484. SRA ids for individual samples can be found in Table S1. A GitHub repository with code can be found at https://github.com/Jcbnunez/Cville-Seasonality-2016-2019. Data S1 and S2 can be found in Zenodo (https://zenodo.org/) at https://doi.org/10.5281/zenodo.7305043. Data S3 can be found in the GitHub repository.

## Supplementary Figures

**Figure S1:**
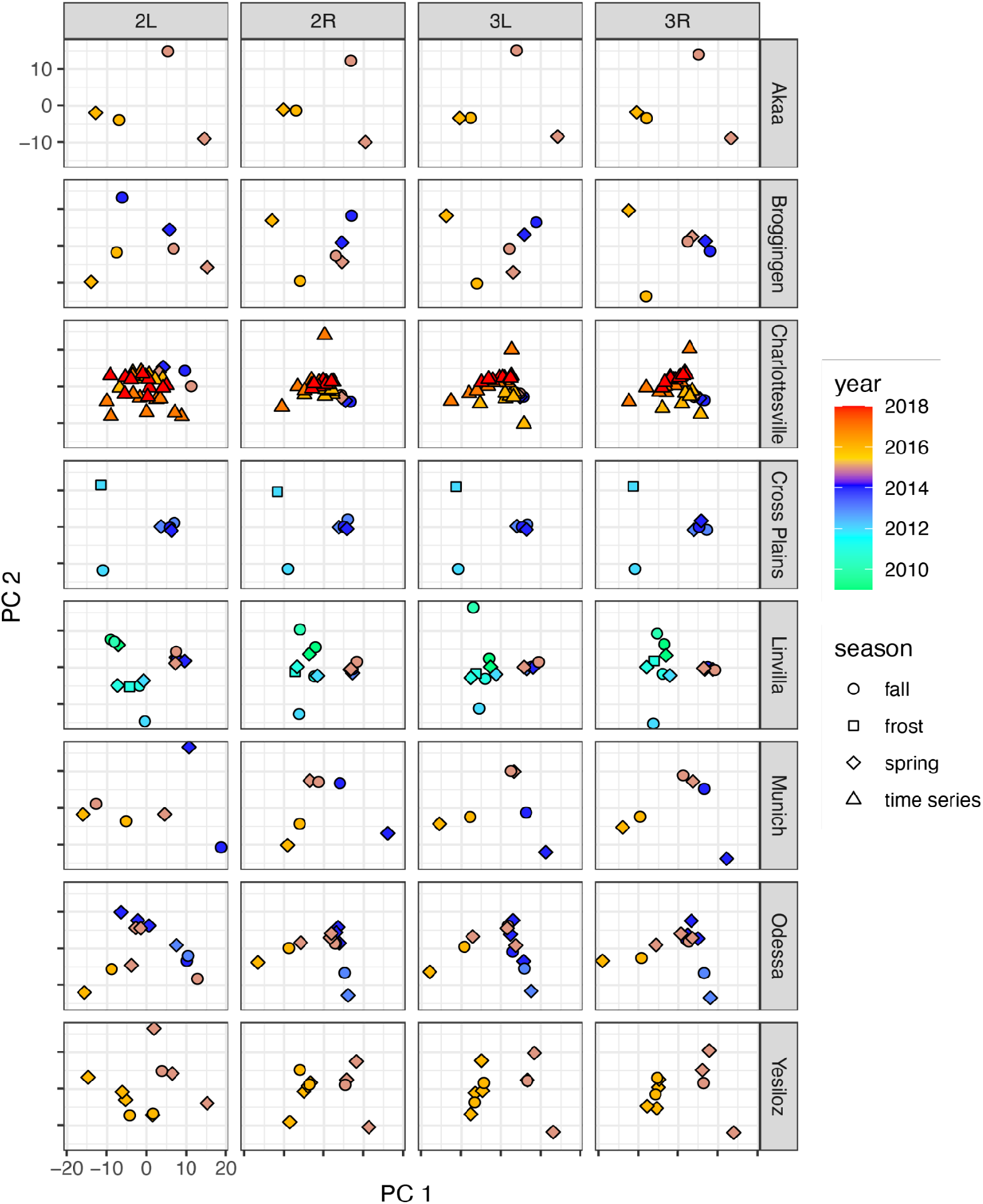
Principal component analysis (PCs 1 and 2 shown), at the chromosome arm level, for each population. The color indicates the collection year. Shapes indicate the sample type: spring and fall collections (collected at the beginning and end of the growing season), frost (collected prior to a frost event), or time series (indicated bi-weekly collections). Samples are: Akaa, Finland; Broggingen, Germany; Charlottesville, VA, USA; Cross Plains, NY, USA; Linvilla PA, USA; Munich, Germany; Odessa, Ukraine; Yesiloz, Türkiye.

**Figure S2:**
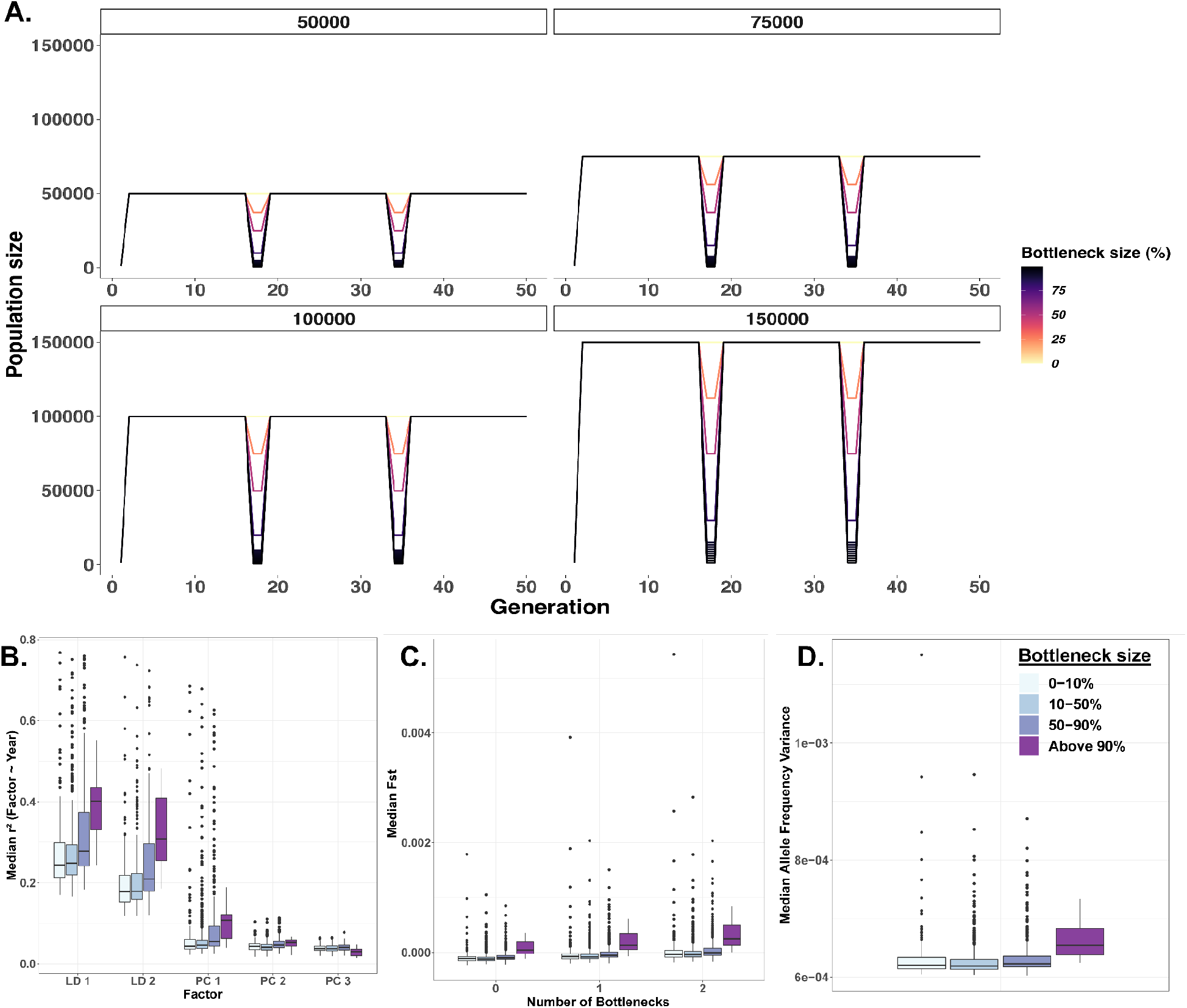
Genetic summary statistics of boom-and-bust simulations. (**A**) Cartoon models of simulated overwintering demography. Each facet showcases example nMax population sizes. (**B**) Median *r*^*2*^ of principal component (PC) and linear discriminant (LD) axis with simulation year. (**C**) Median pairwise-*F*_*st*_ within and between simulation years. D) Median allele frequency variance for simulated models across the entire simulation period. Panels B-D are grouped and colored according to the percentage of bottleneck simulated.

**Figure S3:**
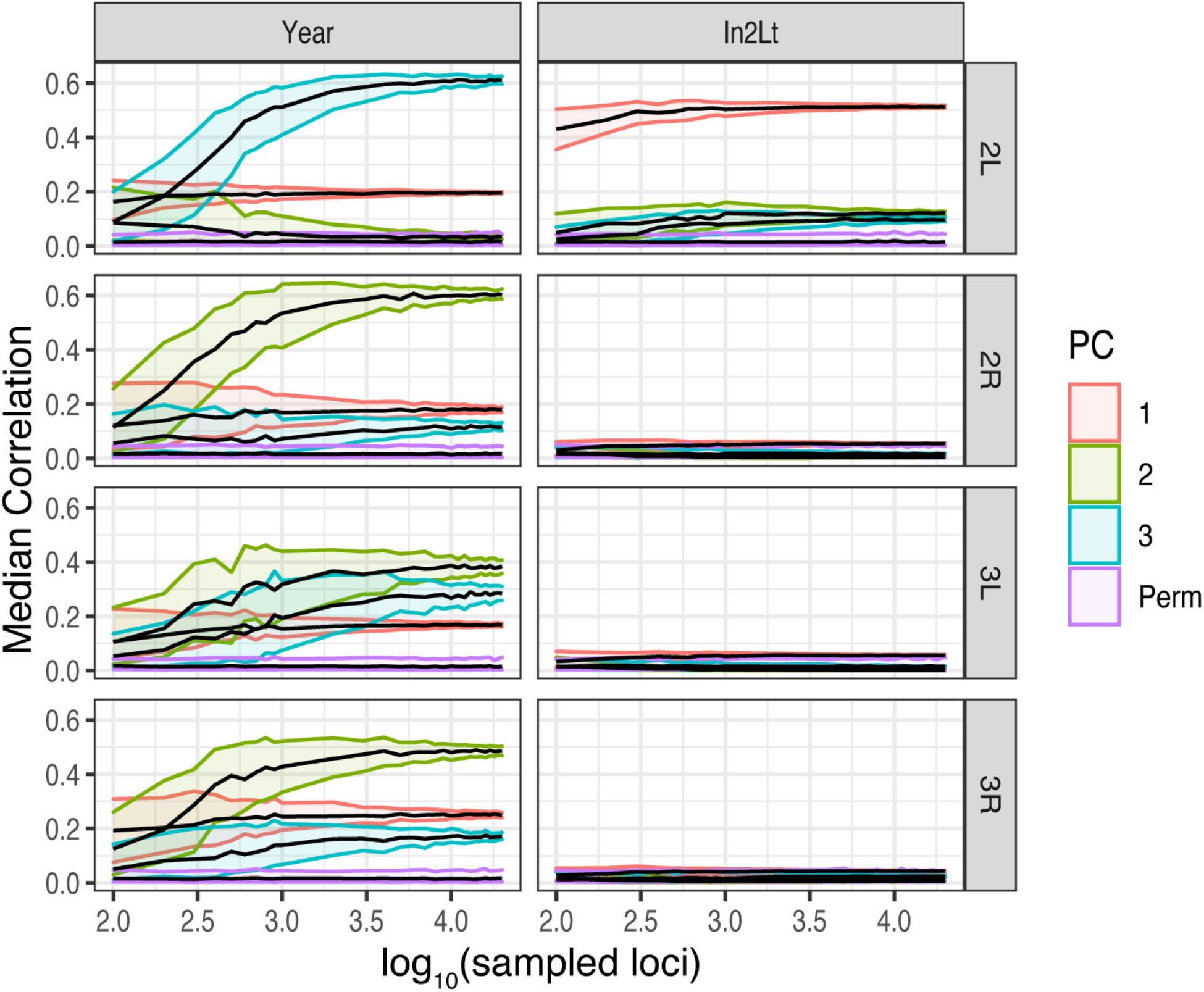
Resampling analysis shows correlations between a number of variables of interest (e.g., Year, Frequency In(2L)t) to the PC projections (PCs 1, 2, and 3) in Charlottesville. Red, green, and blue colors represent PCs 1, 2, and 3 respectively. Permutations are shown in purple.

**Figure S4:**
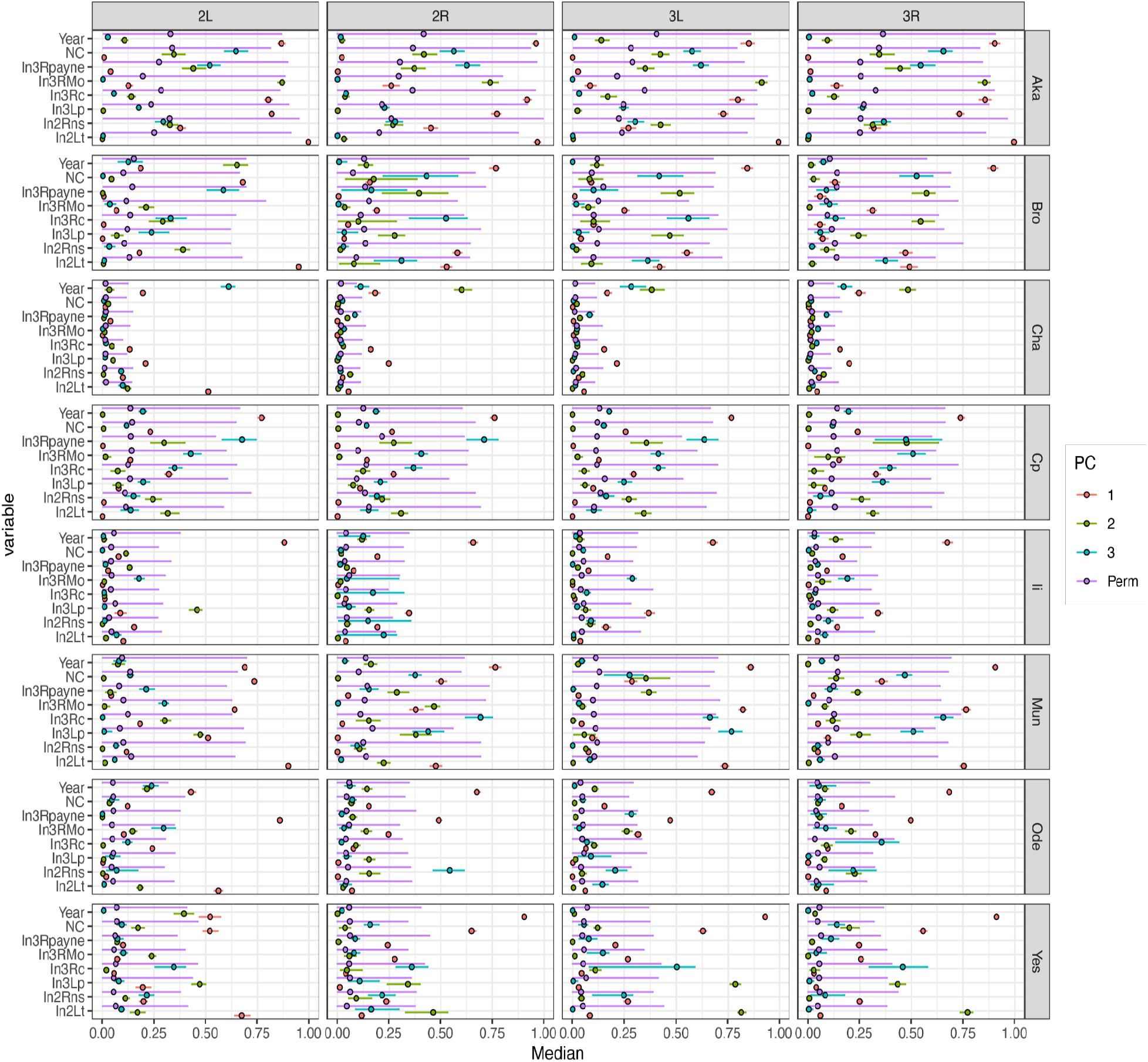
Median correlation between variables of interest (Year, effective coverage, frequency of cosmopolitan inversions) relative to the PC projections (PCs 1, 2, and 3). The confidence intervals represent the 5^th^ and 95^th^ interquartile ranges (IQR). We also show the results of correlations where sample identity has been permuted. Samples are: Aka=Akaa (Finland), Bro=Broggingen (Germany), Cha=Charlottesville (VA), Cp=Cross Plains (NY), Li=Linvilla (PA), Mun=Munich (Germany), Ode=Odessa (Ukraine), Yes=Yesiloz (Türkiye).

**Figure S5:**
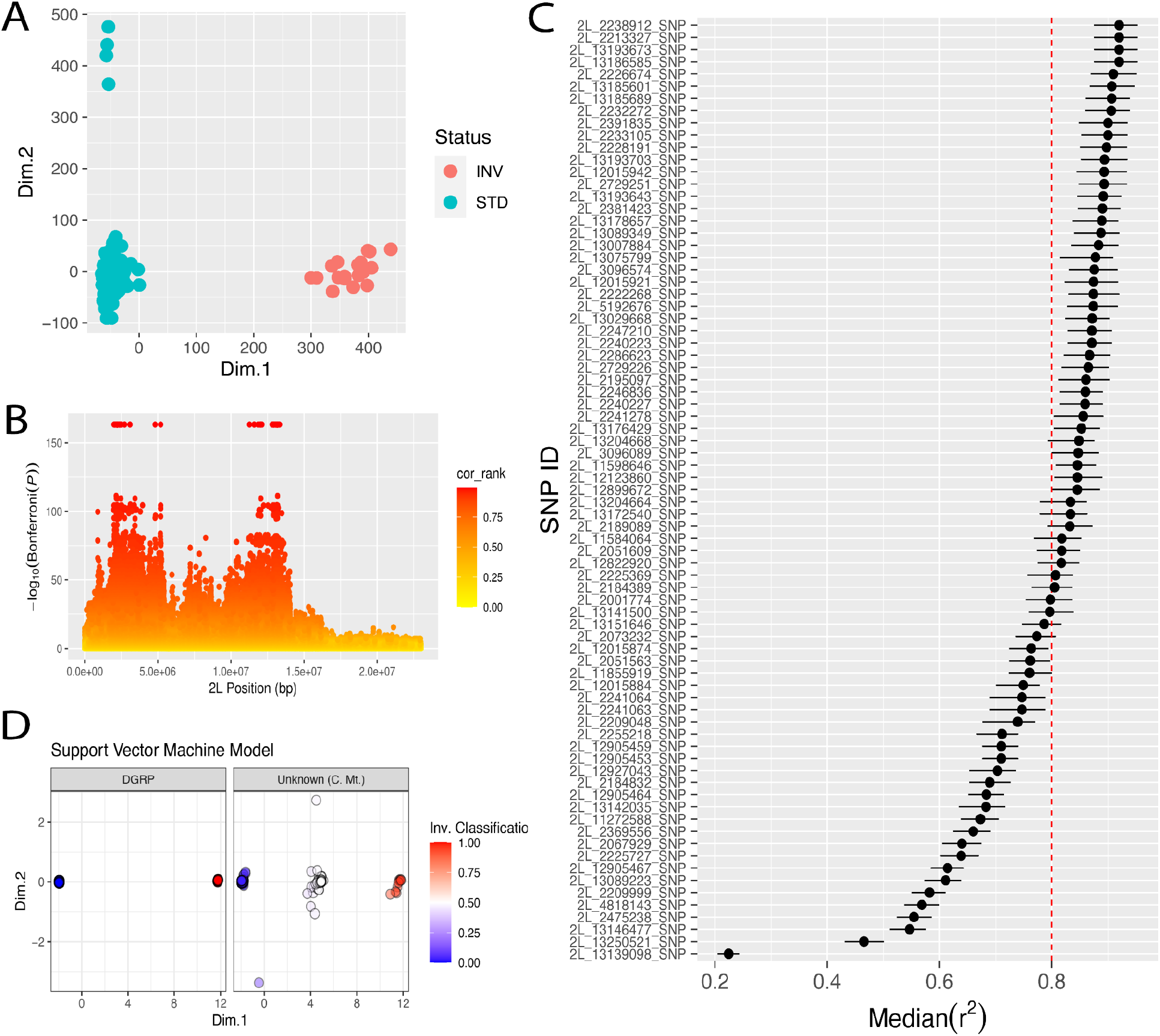
Using the DGRP to characterize mutations associated with the In(2L)t inversion. **(A**) PCA of DGRP lines colored by known inversion status (heterozygous individuals were excluded from the analysis). (**B**) Correlation analysis of individual SNPs within 2L to PC 1 shown in panel A. The y-axis shows the P-value (Bonferroni corrected) of the SNP-level correlation to PC 1 shown in panel A. (**C**) Levels of linkage disequilibrium among the inversion markers discovered in the DGRP estimated in our Charlottesville data. To determine inversion breakpoint markers, we only kept loci with median LD values among markers of 0.8 or greater. (**D**) Results of a support vector machine (SVM) algorithm trained to determine the inversion status of unknown individual samples. The SVM algorithm was trained on the DGRP using the 47 marker loci highlighted on the right side of the dashed red line in C.

**Figure S6:**
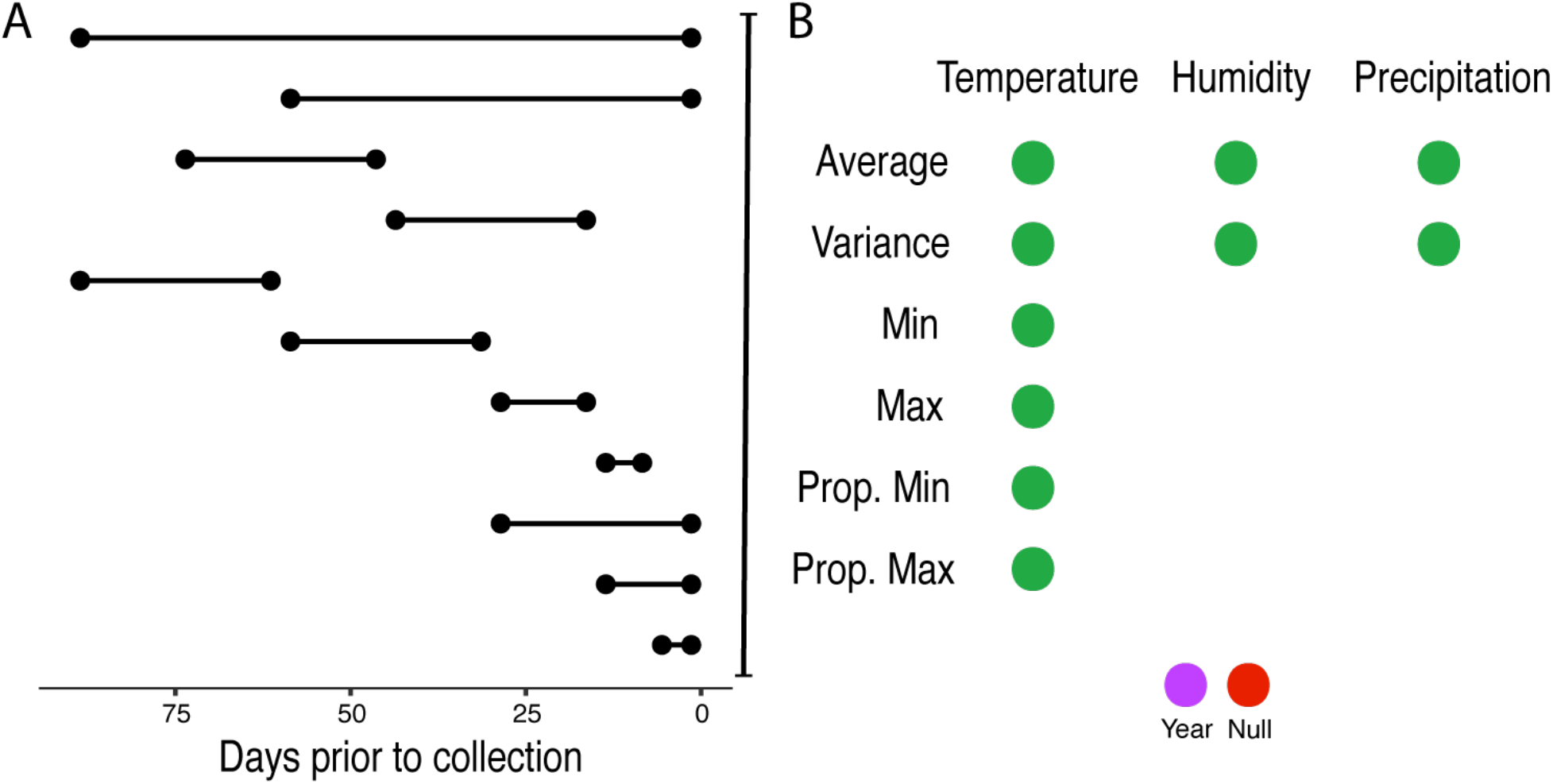
Data summarization scheme using the NASA power dataset for environmental variables. (**A**) The x-axis shows the number of days prior to collection. The panel shows the summary statistics and environmental variables used. (**B**) Green dots indicate that the metric was used. Including a null and year models, the total number of models tested is 11 × 6 (Temperature) + 11 × 2 (Humidity) + 11 × 2 (Precipitation) + 2 (Null and Year) = 112 models.

**Figure S7:**
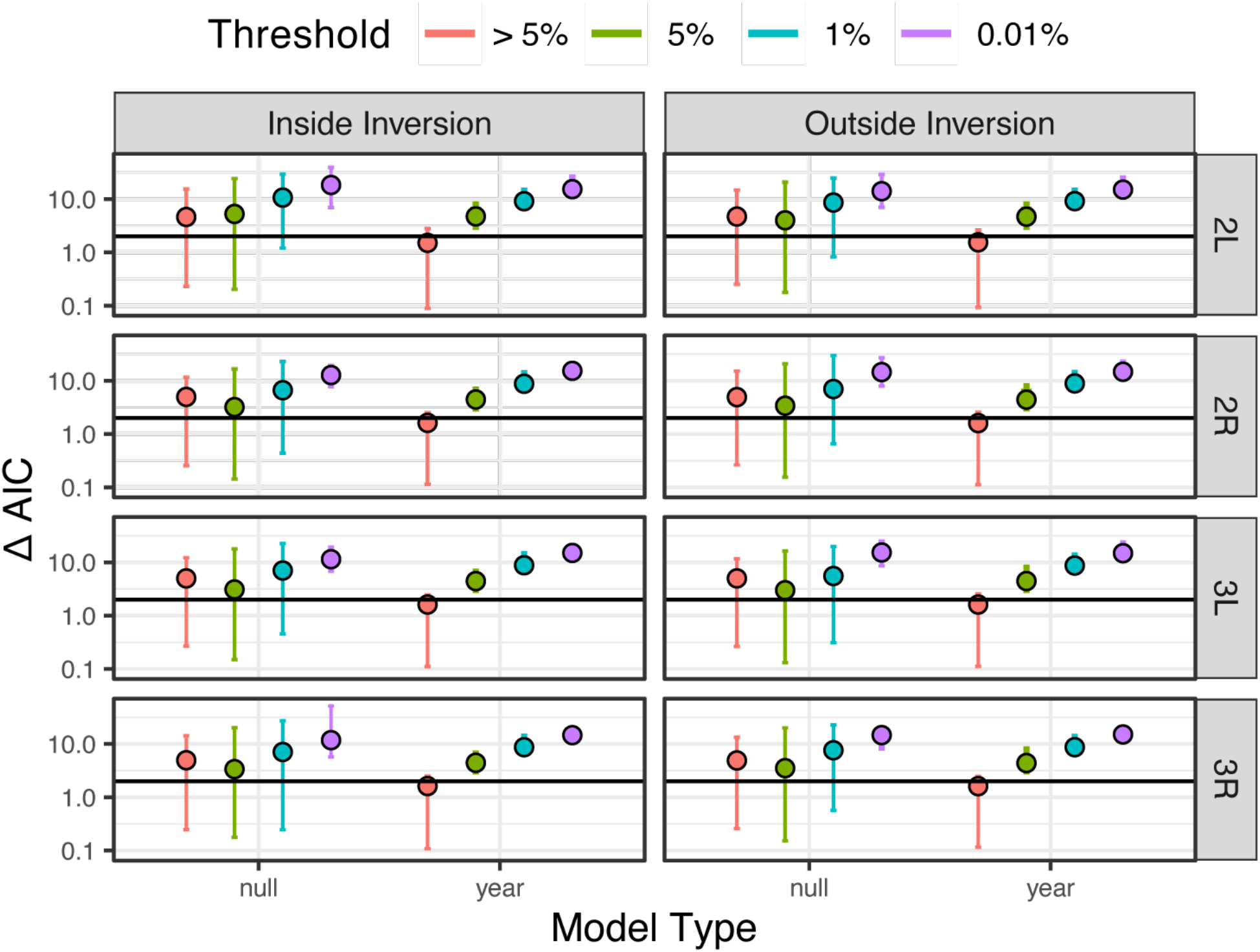
The plot shows ΔAIC that compares various models used in our GLM analyses of Charlottesville. Two comparisons are: 1) the best model in Charlottesville (AF = β_0_ + β_1_(year) + β_2_(Tmax_0-15d_) + *ε*) relative to the null model (AF = β_0_), and 2) the best model in Charlottesville (AF = β_0_ + β_1_(year) + β_2_(Tmax_0-15d_) + *ε*) and the year model alone (AF = β_0_ + β_1_(year)). The horizontal line signifies ΔAIC = 2, a common threshold used for model selection when choosing among models by AIC. The value reported is the median as well as the 2.5% and 97.5% percentiles of ΔAIC for all SNPs across the genome. The color indicates the 3 thresholds in the ranked normalized *P*-value analysis of the GLM.

**Figure S8:**
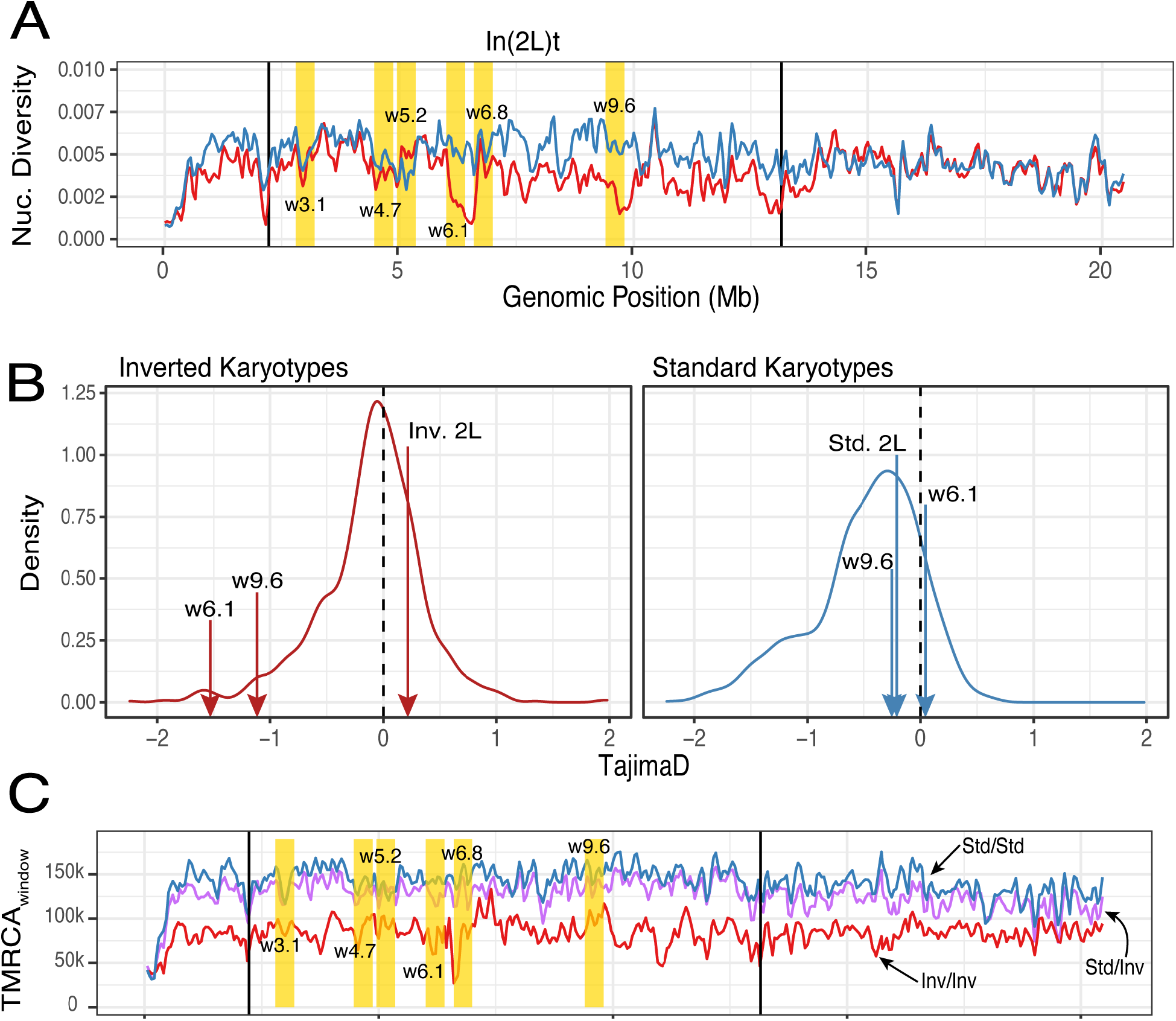
(**A**) Levels of nucleotide diversity (*π*) within chromosome arm 2L. (**B**) Distributions of Tajima’s D for standard and inverted karyotypes. The annotation “Inv” and “Std” refer to the karyotypes In(2L)t and standard, respectively, in chromosome arm 2L. The values for two windows of interest (6.1 and 9.6) are annotated as arrows, as well as the average value for the entire karyotype. (**C**) Distribution of the allele age, defined as time to most common ancestor (TMRCA) across genotypes of Virginian flies. The metric was estimated for standard homozygous (std/std), In(2L)t homozygous (Inv/Inv) and heterozygous (Std/Inv).

**Figure S9:**
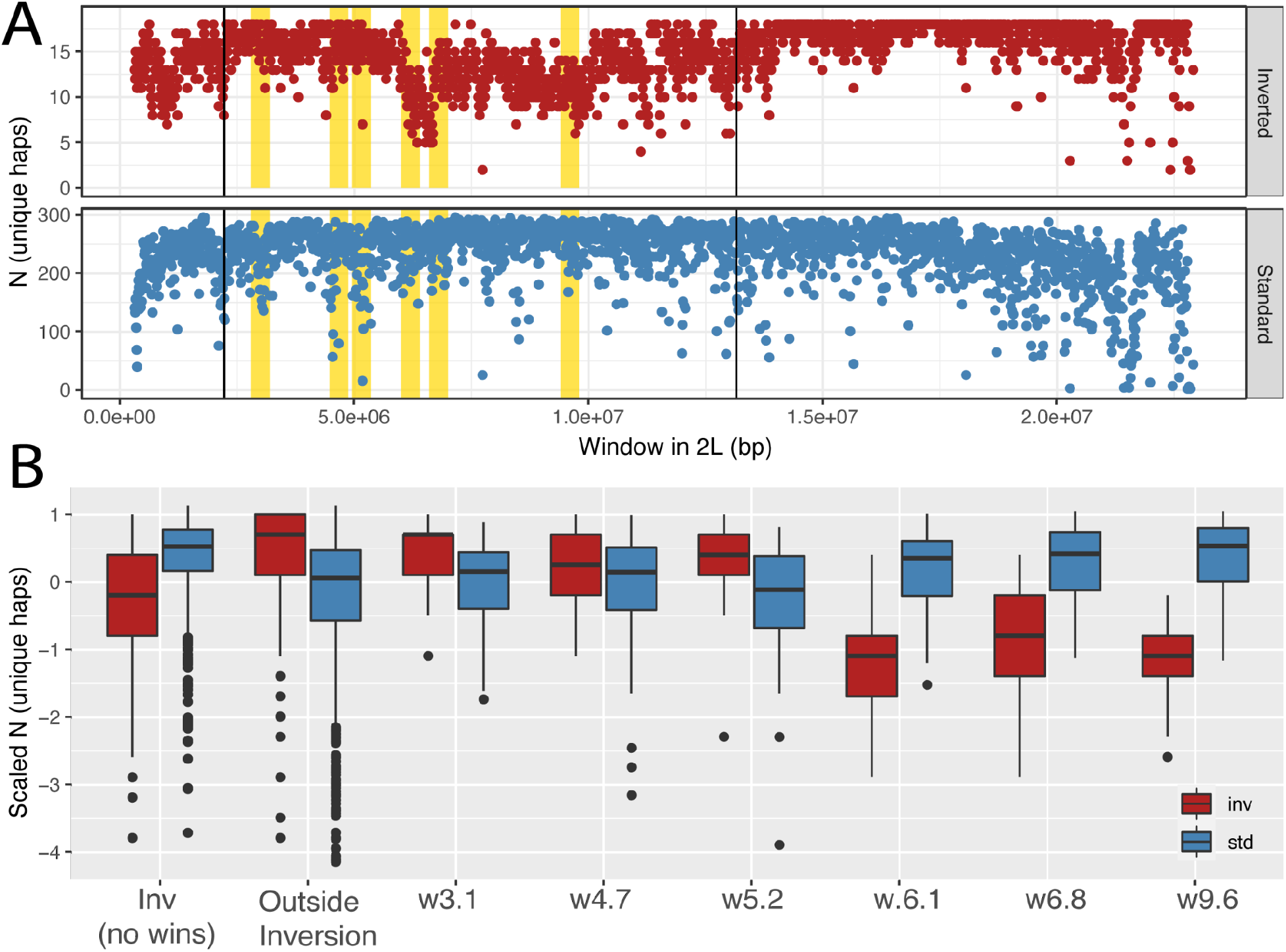
(**A**) Levels of haplotype diversity within inverted and standard classes in chromosome 2L. The y-axis is the number of unique haplotypes across windows of 10k bp. (**B**) Boxplots showing the number of unique haplotypes. The y-axis shows the scaled number of haplotypes (scaled as 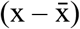; where x: number of haplotypes across individual 10k windows, 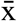 : mean within the region among 10k windows of interest, s: SD within the region of interest). Summaries are shown in six regions of interest in In(2L)t, as well as regions inside and outside of the inversion. The annotation “Inv” and “Std” refer to the karyotypes In(2L)t and standard, respectively, in chromosome arm 2L.

**Figure S10:**
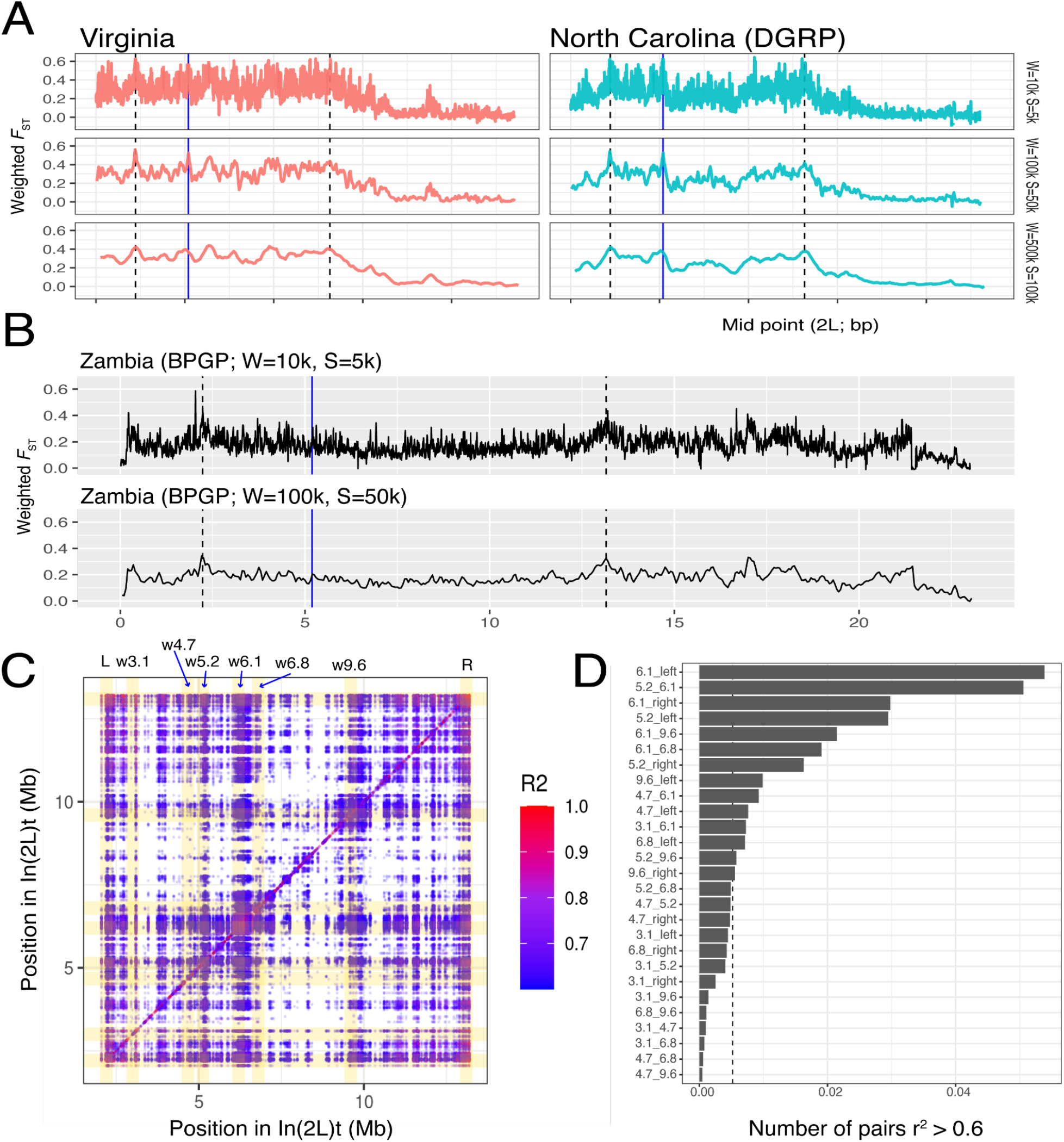
(**A**) *F*_ST_ between the inverted and standard karyotypes in 2L for the Virginia data and the DGRP (North Carolina), both North American samples. The line in blue indicates the trans-species mutation in *Msp300*. (**B**) *F*_ST_ between the inverted and standard karyotypes in 2L in the DPGP (Zambia) an African population. Various window (W) and step sizes (S) are shown. (**C**) Matrix of linkage disequilibrium values (r^2^) within the region of 2L corresponding to In(2L)t. (**D**) Distribution of counts for the number of SNP pairs with r^2^ >0.6 among windows of interest.

**Figure S11:**
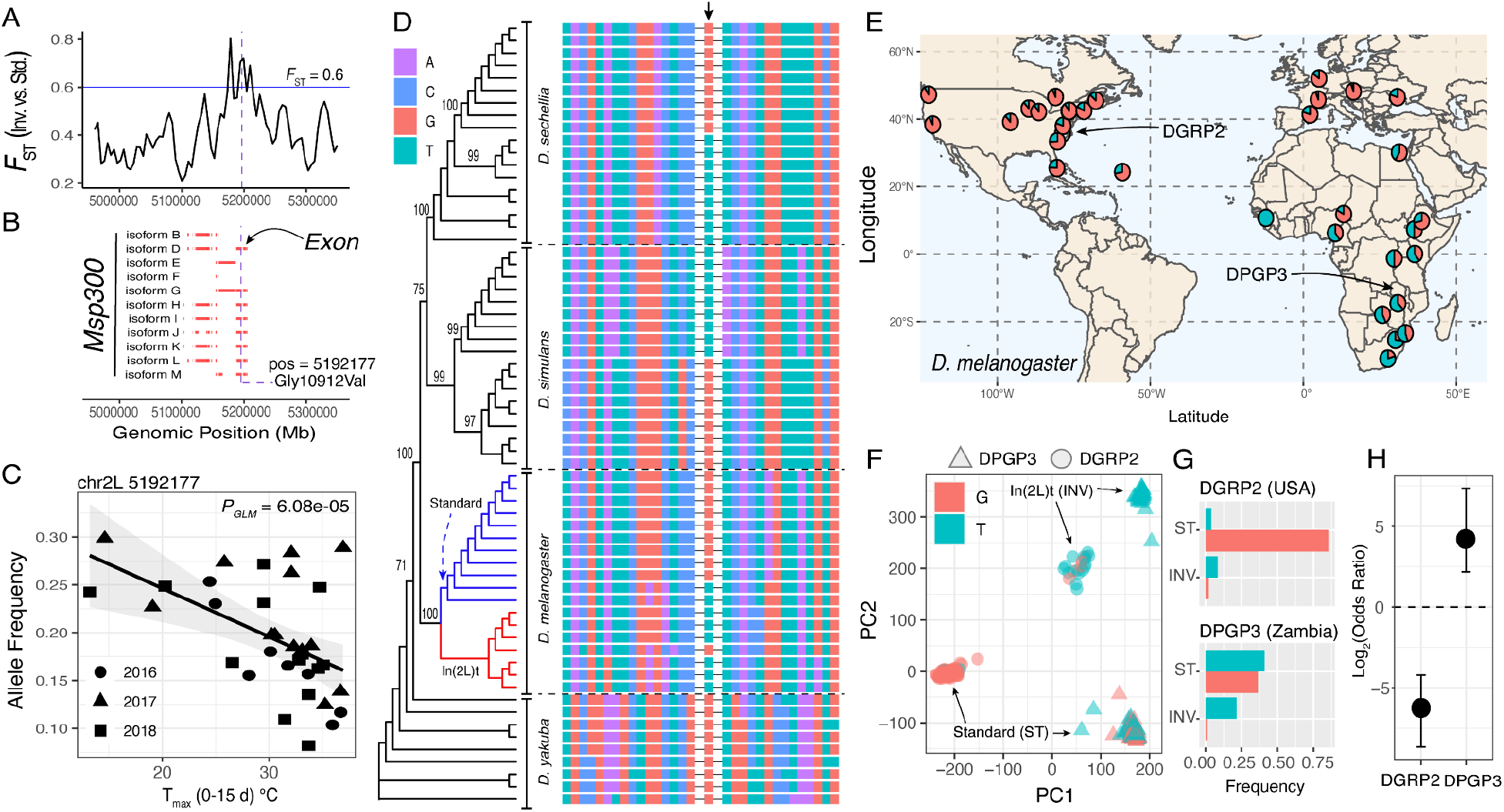
Trans-species polymorphism in *Msp300* drive divergence in 2L. (**A**) Mean *F*_ST_ between inverted and standard karyotype classes, at the *Msp300* region in 2L (Window = 5000 bp, Step = 1000 bp). (**B**) Gene structure and isoforms of *Msp300*. The vertical line indicates the focal mutation (32735G>T, Gly10912Val). (**C**) Correlation between T_max_ (0-15 d) and the frequency of 32735G>T, 32735G>T. (**D**) Phylogenetic tree showing the trans-species polymorphism at 32735G>T (*D. melanogaster* samples from North America are shown). Bootstrap values shown in the nodes. (**E**) Allele frequencies of 32735G>T across populations of *D. melanogaster*. The location of two reference panels, DGRP and DPGP, are shown. (**F**) PCA on temperate (DGRP) and African (DPGP) genetic panels of the locus shows that PC1 mostly captures continental differences between panels. PC2, on the other hand, captures karyotypic differences (For PC1 DGRP/DPGP: *F*_*1,371*_ = 5155.80, *P* = 2.2e-16; For PC1 Inversion: *F*_*1,371*_ = 219.20, *P* = 2.2e-16; For PC2 DGRP/DPGP: *F*_*1,371*_ = 56.10, *P* = 4.885e-13; For PC2 Inversion: *F*_*1,371*_ = 3163.30, *P* = 2.2e-16). (**G**) Frequency of inverted or standard karyotypes carrying 32735G>T in the DGRP and DPGP. (**H**) Enrichment score (Odds Ratio) of T alleles (of 32735G>T) in the standard karyotypes using Fisher’s exact test in the DGRP and DPGP. Notably, patterns of allelic variation are different in African populations. For example, the frequency of T is more abundant in Africa, relative to G (FET, Odds Ratio = 18.43 [95% C.I. = 4.51-16.6]; *P-value* = 3.288e-08) and, unlike temperate populations, both alleles are abundant in the standard karyotype

**Figure S12:**
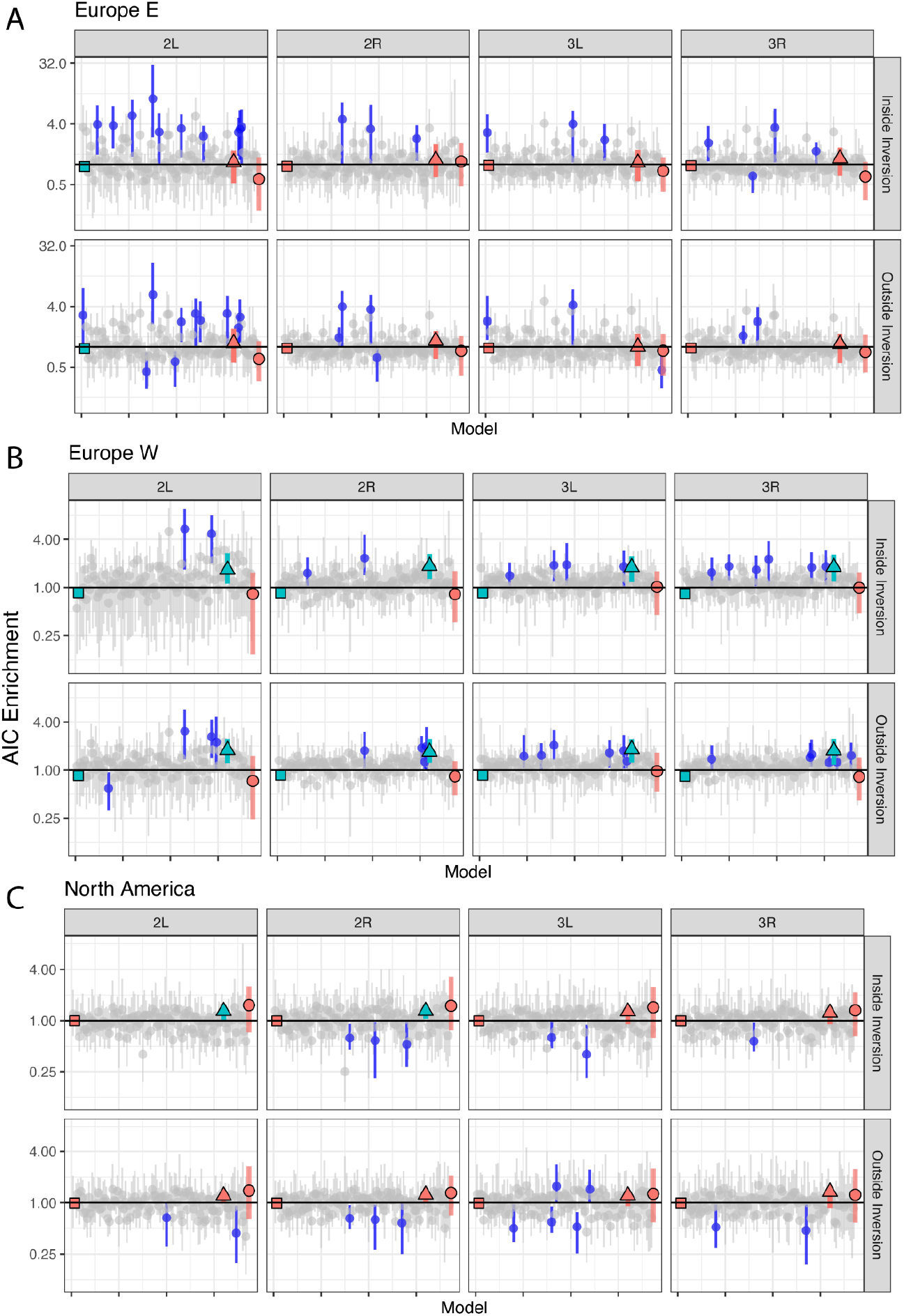
Model search in EU-E (**A**), EU-W (**B**), and NoA-E (**C**). The x-axis shows each model ranked according to the best model in In(2L)t in Virginia. The y-axis shows an enrichment score testing whether the model output was scored as the best model by AIC relative to permutations. The vertical line represents the expected value across all permutations. Confidence intervals are reported as the 1% and 99% percentiles of the estimators. Gray circles represent models that are not statistically significant. Blue circles represent models that are statistically significant. There are three special models highlighted in this plot. First: T_max_(0-15 d) is shown as a black circle that is always the first model (since it is the best model in Virginia). Second: the year model shown as a triangle. Third: the null model shown as a square. For these three special models, green indicates significance whereas red the lack of significance.

**Figure S13:**
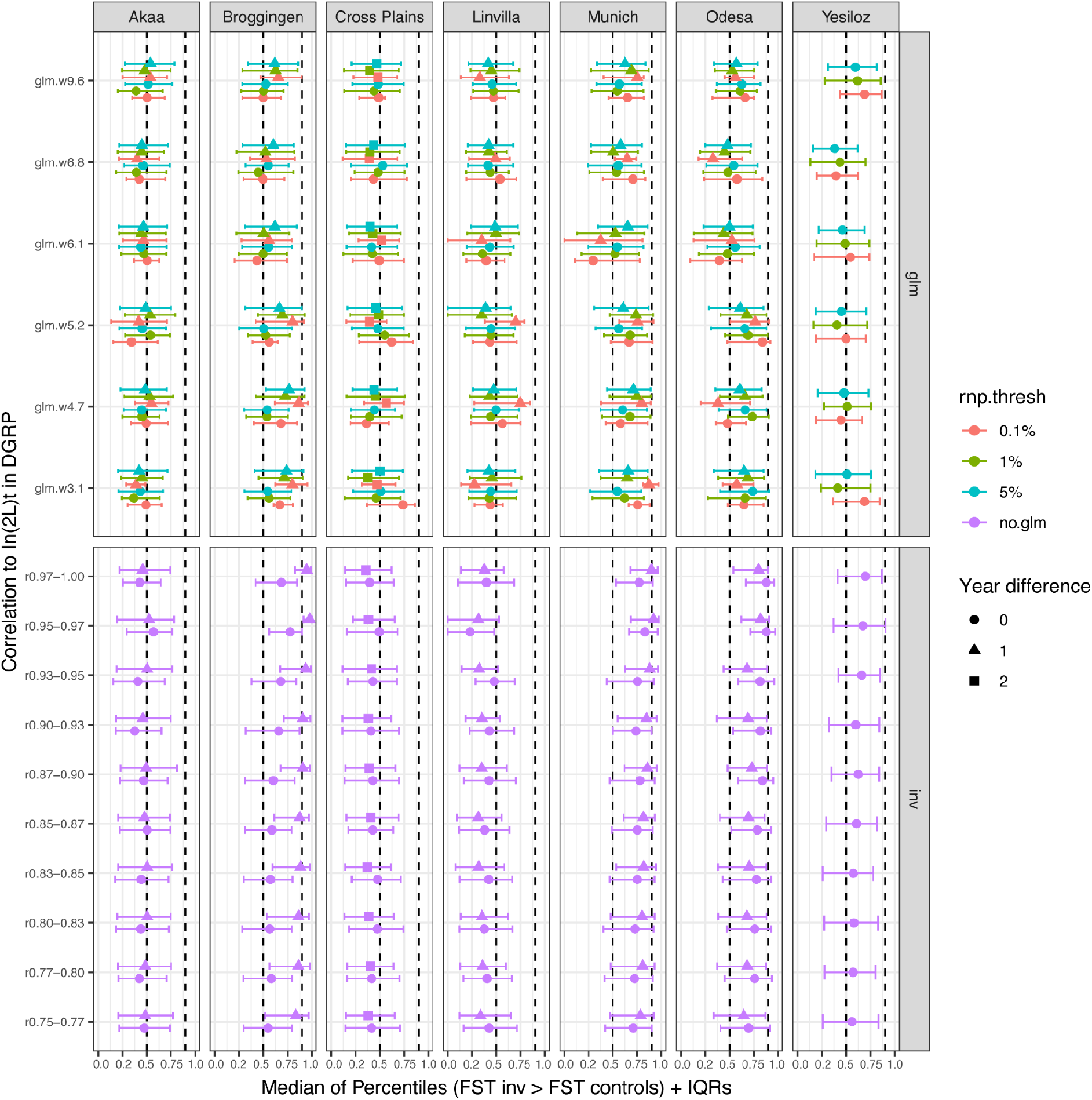
Mean of the percentile of *F*_ST_ values from loci associated with In(2L)t relative to matched controls for various locales. The top panel shows windows of interest in the GLM analysis. The lower panel shows the correlation rank of loci correlated to the inversion (as per PCA analysis of DGRP lines). The x-axis shows the percentile of control loci with *F*_ST_ values lower than that of the locus of interest. Interquartile ranges (IQRs) are shown.

**Figure S14:**
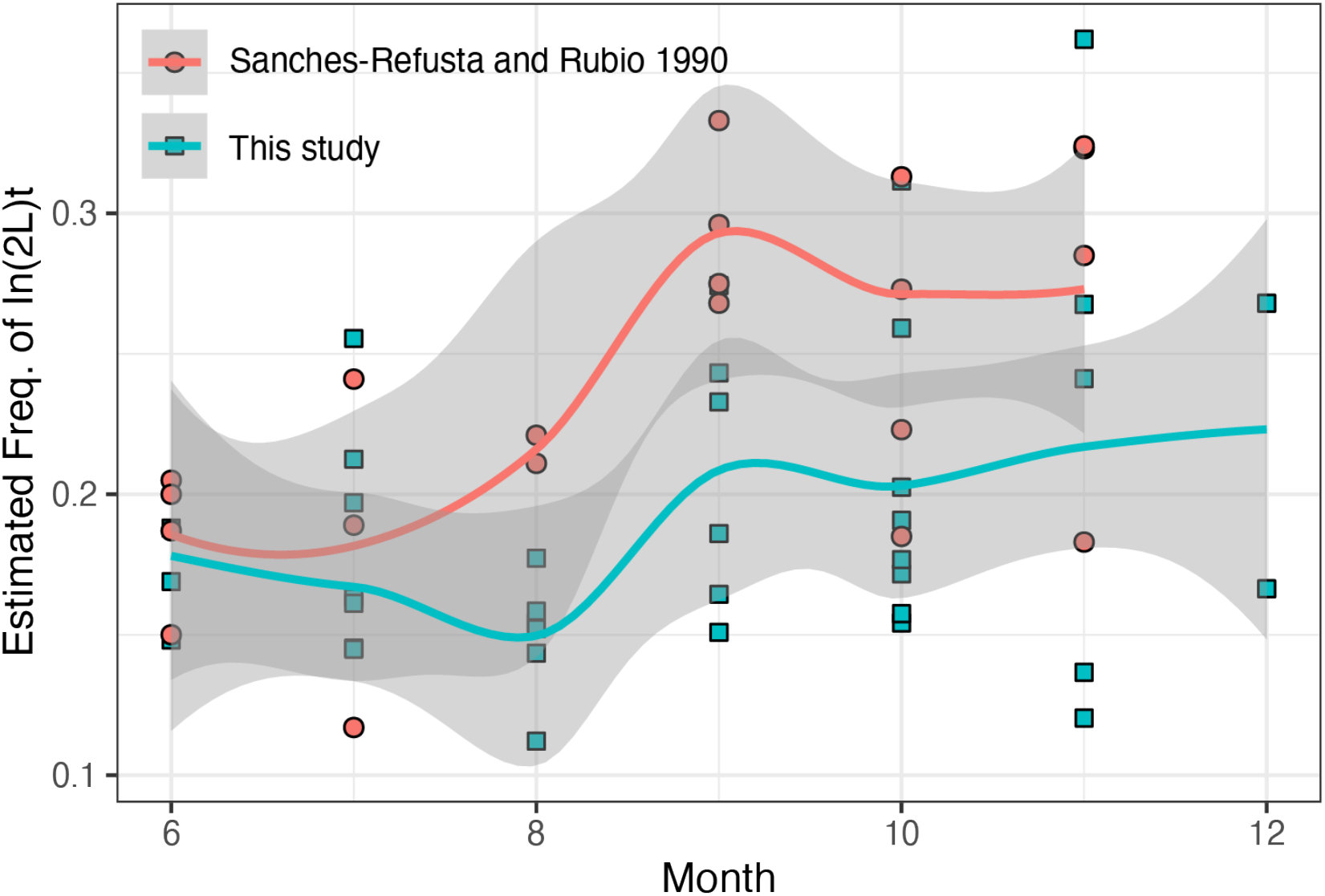
Estimated frequencies of the inversion In(2L)t across a previous study and this one looking at changes in response to seasonality. The month-to-month correlation between the patterns of allele frequency between this paper and our study is 0.76 (*P* = 0.003).

**Figure S15:**
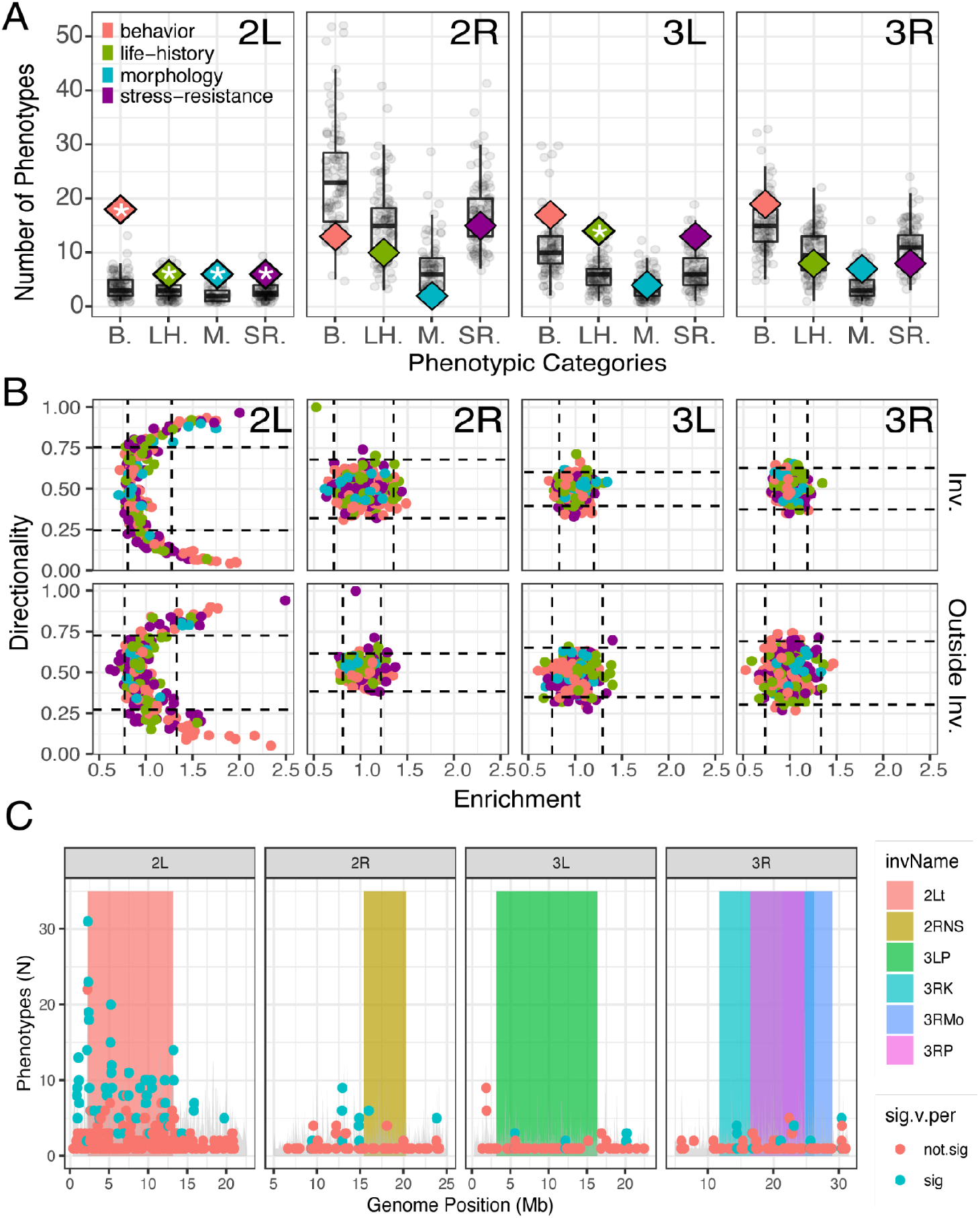
**(A)** The y-axis shows the number of traits used in GWAS that are associated with inversion status in the DGRP for autosomes (For chromosome 3R the test considers whether a trait is associated with any inversion). Traits are divided across four phenotypic categories indicated by color. The real data is shown as diamonds, permutations are shown as black points and boxplots (asterisks indicate that the data outperform 95% of permutations). (**B)** Directionality and enrichment analysis between the DGRP-GWAS and the best environmental model in Virginia for 2R, 3L, and 3R. Lines indicate the permutation based 95% confidence intervals. (**C)** Window level enrichment analysis across the whole genome. Windows that beat permutation are shown in turquoise, otherwise in red.The y-axis shows the SNP-wise number of enriched phenotypes (i.e., significant in both the GLM and GWAS). Inversions are shown as color blocks.

**Figure S16:**
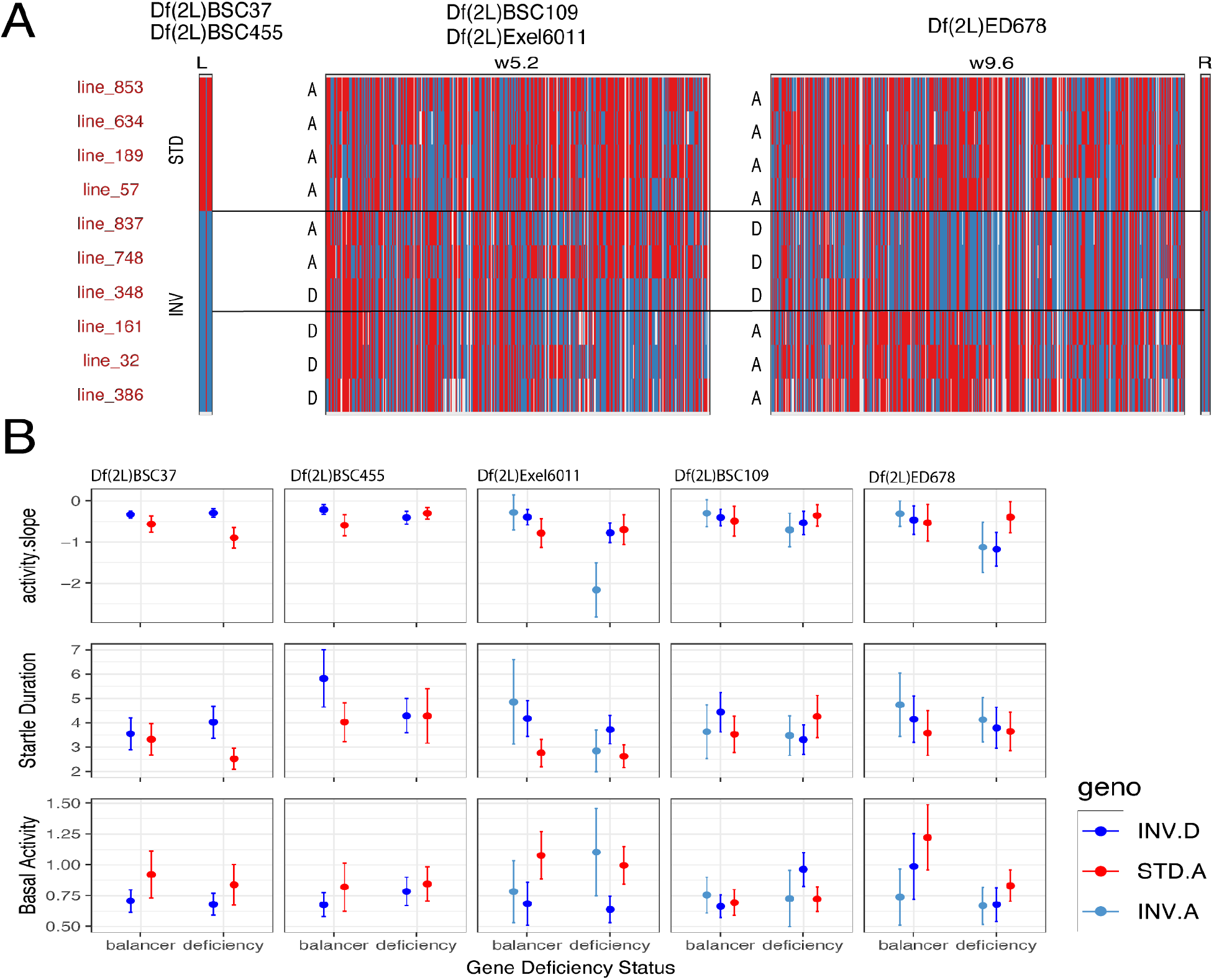
Explanation of DGRP line choice and the complete observations of three major phenotypes from each the different genetic backgrounds used in the deficiency line startle response study. (**A)** The ancestral (red) or derived (blue) background of the 10 DGRP lines used in the study. Lines are shown to be both inverted or standard, and when considering the structure in their genetic backgrounds either in the “A” type or “D” type (based on the proportion of sites shared or derived relative to *D. simulans*). For example, flies with DGRP 837 background are inverted/type A when considering the 5mb window, and type D when considering the 9mb window. (**B)** The differences in the three observed phenotypes are shown across genetic groups. Phenotypes are faceted by each of the 5 major deficiencies (see **Tables S10, S11**), grouped by the presence of deficiency or balancer in the F_1_ flies used, and colored by their inversion and type. The top row illustrates the activity slope, a measurement of how rapidly activity decays back to basal following the stimulus event. The second row illustrates the startle duration, a measurement of the period between the peak of activity following stimulus, and the return to basal activity. Third row indicates the level of basal activity per minute.

## Supplementary Tables (Legends)

**Table S1:** Samples used in this analysis. The table has the following headings: sampleId: the name of the pooled or individual sample; country: country of provenance; city: city of provenance; locality: code indicating the country, state, and city where the sample was collected; collectionDate: collection date; nFlies: number of flies pooled, for individual samples this number is 1; SRA_accession; SRA_experiment; type: pooled or individual; PCA.set: TRUE/FALSE whether the sample was used in the PCA analysis; FST.set: TRUE/FALSE whether the sample was used in the *F*_ST_ analysis; GLM.set: TRUE/FALSE whether the sample was used in the GLM analysis; IND.set: TRUE/FALSE whether the sample was used in the individual analyses. in(2L)t_score: For pooled samples, this value is the frequency of the inversion In(2L)t in the pool. For individual samples, this is the SVM score for a sample having inverted (1), standard (0), or heterozygous (0.5). propSimNorm: The level, standardized, of *D. simulans* contamination in pooled data. Note that metadata for DEST samples can be found at: https://github.com/DEST-bio/DEST_freeze1/tree/main/populationInfo.

**Table S2:** Correlation analysis between PC 1 projections for each city and the year of collection of the pool. Headers: City: city of provenance; Country/State: country or state from which samples originated; PC: Principal component used to run the correlation; r: The correlation between the collection year and the PC projection; r^2^: The coefficient of determination of the model; P-value: P-value of the model; Sig: whether the P-value is below 5%.

**Table S3:** *P-values* for the population-level Kruskal–Wallis (Overwintering test).

**Table S4:** Full results for resampling analysis of correlations between PC projections and several variables across populations. Columns shown are: Population, chromosome, number of SNPs sampled, Principal Component, Median correlation, IQR25, IQR75, IQR05, IQR95, Mean, SD.

**Table S5:** Tukey HSD analysis on the haplotype analysis of In(2L)t. Columns shown are: comparison (com), difference in means (diff), lower 95% confidence interval (lwr), upper 95% confidence interval (upr), *P-value* adjusted (p.adj).

**Table S6:** Metadata for additional single individuals from North Carolina, Pennsylvania, Maine, France, and the Netherlands. Columns are: Sample name, location, longitude, latitude, Citation where the data can be found.

**Table S7:** Output of the SVM prediction model. Columns: sample Id, quantitative prediction from the SVM, qualitative prediction from the SVM, population.

**Table S8:** Phenotypes used in our meta-analysis including their references and DOIs. Columns: id, phenotype name, doi of study, specific phenotype category, general phenotype category.

**Table S9:** Line averages for phenotypes in the DGRP.

**Table S10:** Deficiency lines and DGRP crossing scheme.

**Table S11:** Statistical output of complementation tests comparing different phenotypes across the 5 different deficiencies backgrounds.

## Supplementary Datasets (Legends)

**Data S1:** Data object including the SNP-wise GLM output. The columns in this dataset are as follows: chr (chromosome), pos (position), AIC (AIC of the model at given SNP), variable (environmental variable from NASA-POWER), mod_id (Model Id), p_lrt (LRT p value), b_temp (beta of environemtnal variable), se_temp (SE of environemtnal variable), Cluster (Population, i.e., VA, EUE, EUW), cm_mb (Recombiantion), invName (Inversion Name), rnp (Ranked Normalized P-value), Perm.rnp.0.01.quant (0.01 quantile of RNP in 100 permutations), time_window (window of time of the model), SNP_id (SNP id), N.phenos (Phenotype number), Description (Pheno description)

**Data S2:** Data object including the AIC models enrichment analysis. Headers are: variant.id (variant Id in DESTv1.1), perm (0: real data, 1-100: indicates permutation number), cluster (population cluster), var (ecological variable), mod (summarization scheme), p_lrt (likelyhood ratio test), minAIC (minumun AIC of the ecological variable model), yearAIC (AIC of year model), chr (chromosome), pos (position), N (number repetitive libraries; all should be 0), libs (repetitive elements; all should be filtered), cm_mb (recombination rate), invName (inversion status and name). The following columns indicate whether the SNP passes filters across core populations: DE_Bro, DE_Mun, FI_Aka, PA_li, TR_Yes, UA_Ode,, VA_ch.

**Data S3:** Anchor SNPs and associated pairs in high LD. This data contains the following headers: sampleId (sample Id), variant.id (variant Id in DESTv1.1), ad (allele count), dp (coverage), col (annotation: function), gene (annotation: gene name), chr (chromosome), pos (position), nAlleles (allele number), af (allele frequency), country, city, collectionDate, lat, long, season, locality, type, continent, set (population set), nFlies (number of flies pooled), SRA_accession, SRA_experiment, SeqPlatform (sequencing platform), collector, sampleType, year, yday (day of collection), stationId (weather station info), dist_km, nEff (effective coverage), af_nEff, win (window of interest in In2Lt), sim_af (Simulans allele), af_polarized (allele frequency polarized to simulans).

